# Genome-wide association mapping of ethanol sensitivity in the Diversity Outbred mouse population

**DOI:** 10.1101/2021.09.06.459155

**Authors:** Clarissa C. Parker, Vivek M. Philip, Daniel M. Gatti, Steven Kasparek, Andrew M. Kreuzman, Lauren Kuffler, Benjamin Mansky, Sophie Masneuf, Kayvon Sharif, Erica Sluys, Dominik Taterra, Walter M. Taylor, Mary Thomas, Oksana Polesskaya, Abraham A. Palmer, Andrew Holmes, Elissa J. Chesler

## Abstract

**Background:** A strong predictor for the development of alcohol use disorders (AUDs) is altered sensitivity to the intoxicating effects of alcohol. Individual differences in the initial sensitivity to alcohol are controlled in part by genetic factors. Mice offer a powerful tool for elucidating the genetic basis of behavioral and physiological traits relevant to AUDs; but conventional experimental crosses have only been able to identify large chromosomal regions rather than specific genes. Genetically diverse, highly recombinant mouse populations allow for the opportunity to observe a wider range of phenotypic variation, offer greater mapping precision, and thus increase the potential for efficient gene identification.

**Methods:** We have taken advantage of the Diversity Outbred (DO) mouse population to identify and precisely map quantitative trait loci (QTL) associated with ethanol sensitivity. We phenotyped 798 male J:DO mice for three measures of ethanol sensitivity: ataxia, hypothermia, and loss of the righting response. We used high density MEGAMuga and GIGAMuga arrays to obtain genotypes ranging from 77,808 – 143,259 SNPs. In addition, we performed RNA sequencing in striatum to map expression QTLs and to identify gene expression-trait correlations.

**Results:** We then applied a systems genetic strategy to identify narrow QTLs and construct the network of correlations that exist between DNA sequence, gene expression values and ethanol-related phenotypes to prioritize our list of positional candidate genes.

**Conclusions:** Our results can be used to identify alleles that contribute to AUDs in humans, elucidate causative biological mechanisms, or assist in the development of novel therapeutic interventions.

## Introduction

Alcohol use disorder (AUD) is a common psychiatric disorder with a substantial genetic component (Kendler et al. 2012; Verhulst et al. 2015). In recent years, genome-wide association studies (GWAS) have identified genetic variants associated with AUD and self-reported problematic alcohol use (Gelernter et al. 2014; Quillen et al. 2014; Mbarek et al. 2015a; Kranzler et al. 2019; Sanchez-Roige et al. 2019a, b; Zhou et al. 2020). However, human GWAS also face a number of challenges, including high costs associated with phenotyping and genotyping such extremely large samples, typical use of binary traits that lack precision, reliance on self-report for many relevant phenotypes, a genetic architecture in which most SNPs have very small effect sizes, the presence of rare alleles/low minor allele frequencies, limited access to relevant tissues for gene expression analysis, and difficulties associated with functional validation studies (Hindorff et al. 2009; Stranger et al. 2011; Ward and Kellis 2012; Hart and Kranzler 2015).

An alternative approach now gaining traction in large-scale human AUD GWAS involves endophenotypes such as alcohol consumption, electroencephalogram oscillatory activity, and most recently, sensitivity to the effects of alcohol (Meyers et al. 2016, 2017, 2020; Liu et al. 2019; Gelernter et al. 2019a; Lai et al. 2020b, a; Sanchez-Roige and Palmer 2020). In humans, there is dramatic individual variability in sensitivity to both the positive and negative effects of alcohol. This variability is partly genetic in origin (Grant 1998; Schuckit and Smith 2000, 2001), has been associated with subsequent risk for development of AUD (Newlin and Thomson 1990a; Schuckit 1994; Holdstock et al. 2000; Krystal et al. 2003; Schuckit and Smith 2011; King et al. 2011; Schuckit et al. 2014), and is predictive of treatment success (Savage et al. 2015; Schuckit et al. 2016). Therefore, the acute subjective effects of alcohol may be a clinically relevant endophenotype for AUD. However, traditional laboratory-based alcohol challenges are expensive and time-consuming, making them impractical to use at the scale needed for GWAS. As a result, they have not been widely utilized in human GWAS, even though they have been of great interest to the alcohol research community for many years (Quinn and Fromme 2011).

One AUD endophenotype that can be readily assessed in mouse models is ethanol sensitivity (Parker et al. 2020). As in humans, ethanol sensitivity in mice is determined partly by genetic factors (Crabbe et al. 1989, 1994; Christensen et al. 1996; Markel et al. 1997; Gehle and Erwin 2000; Chesler et al. 2012; DuBose et al. 2013). However, traditional mouse mapping studies used populations that typically lacked sufficient amounts of recombination that were needed for gene identification (Parker and Palmer 2011; Saul et al. 2019).

This limitation can be overcome by using populations with more accumulated recombinations to improve mapping resolution. The mice used in the present study are Diversity Outbred (DO) stock. DO mice are a unique resource that provide the advantages of working with a mouse model while still maintaining large amounts of recombination that more closely approximate that of a human GWAS. The DO population is a genetically and phenotypically heterogeneous population derived from five classical (A/J, C57BL/6J, 129S1/SvlmJ, NOD/ShiLtJ, NZO/H1LtJ) and three wild-derived (CAST/EiJ, PWK/PhJ, WSB/EiJ) mouse strains (Churchill et al. 2012). DO mice have an elevated minor allele frequency across the genome and levels of recombination favorable for high-resolution genetic mapping (Svenson et al. 2012). Furthermore, the DO population is extendable: each mouse is genetically unique and additional cohorts can be added to studies as they proceed (Chesler et al. 2014).

By using mice as our model organism, we can employ pharmacological challenge assays reminiscent of laboratory-based alcohol challenges in humans, but in a high-throughput manner using smaller sample sizes as compared to human GWAS that still provide sufficient power for gene discovery. Furthermore, we can use alcohol naïve subjects for studies of sensitivity, thus removing important confounding factors that are unavoidable even in the best human laboratory studies of alcohol sensitivity. In the present experiment, we characterized a DO mouse population on three measures of ethanol sensitivity, genotyped them using next-generation genotyping arrays, and performed genetic linkage and SNP association mapping using updated statistical algorithms. We identified narrow chromosomal regions associated with our traits of interest and used a series of bioinformatic approaches in conjunction with RNA-Seq data to map expression QTLs (eQTLs) and prioritize candidate genes from within these loci for future functional validation.

## Materials and methods

### Animals and housing

All procedures were approved by the NIAAA and Middlebury College IACUC in accordance with NIH guidelines for the care and use of laboratory animals. Male DO mice (N = 798) were obtained from Jackson Laboratory (JAX stock number 009376 (J:DO); Bar Harbor, ME, USA) from outbreeding generations G9, G11, G16, G17, G18, G20, and G21. Mice were phenotyped at either NIAAA (N = 290) or Middlebury College (N = 508). Mice were individually housed in temperature (23 ± 2°C) and humidity (45 ± 15%) controlled vivariums, under a 12-h light/dark cycle (lights on at 0630 NIAAA/0700 Middlebury) with standard laboratory chow (Teklad 2020X, Envigo, Madison, WI, USA) and water available *ad libitum*. All mice were involved in a separate behavioral study of conditioned fear (data not shown), after which (>1 week) they were tested for ethanol sensitivity.

### General procedures

A battery of three assays for intoxication was employed: 1) ethanol-induced ataxia, 2) hypothermia, and 3) loss of the righting response (LORR). For each test, mice were transported from the adjacent vivarium and allowed to habituate to the procedure room for one hour in their home cages. All mice were tested on each assay with the test involving the lowest ethanol dose (i.e., ataxia) first, followed by hypothermia and then LORR, with an interval of at least one week between tests. Long-term tolerance to ethanol’s effects is not expected to occur with this test and treatment regimen (Crabbe et al. 2008). For all assays, ethanol (200 proof, prepared in 0.9% saline to produce 20% v/v solutions) was administered via intraperitoneal (i.p.) injection with dose determined by manipulating the volume of injection. Mice were weighed on day of testing. Temperature of the testing rooms was maintained at 21 ± 2°C. Additional details regarding phenotyping procedures are provided in the supplementary materials.

### Genotyping

Mice were euthanized by cervical dislocation followed by immediate decapitation at ∼164.6 days (SD = 44.9). Euthanasia and dissections were performed between 8AM-2PM. Tail and spleen samples were removed from each animal for genotyping, placed into 1.5mL Eppendorf tubes, and stored in saline at −80°C. Tail samples were shipped to Neogen Inc. (Neogen Inc., Lincoln, NE, USA) for genotyping. NIAAA mice were genotyped on the MegaMUGA (N = 277), and mice tested at Middlebury were genotyped on the GigaMUGA (N = 500) Illumina array platforms. The MegaMUGA assays 77,808 markers spanning the 19 autosomes and X chromosome of the mouse with a mean spacing of 33 Kb. The GigaMUGA improves on the MegaMUGA by increasing the marker density to 143,259 markers (approximately half of which are carried over from the MegaMUGA), with average spacing of 18 Kb (Morgan et al. 2015). Markers were optimized for information content in the Collaborative Cross (CC) and the DO mice.

### RNA extraction and sequencing for expression-trait correlations

In a subset of the Middlebury DO mice that were tested for ethanol sensitivity, whole brains were removed and incubated in a container of chilled RNALater buffer solution (Thermo Fisher Scientific, Waltham, MA, USA) for one minute before dissecting hippocampus and striatum. Individual brain regions were then placed in 1mL of RNALater buffer and immediately stored on ice. Samples were then transferred and stored in a −80°C freezer prior to RNA extraction. Hippocampal and striatal tissue were homogenized, and total RNA was extracted as previously described (Yazdani et al 2015; Bryant et al. 2012), then purified using the RNeasy kit (Qiagen, Valencia, CA, USA). RNA was shipped to the UCSD Genomics Core Facility where libraries were prepared from total RNA with RIN score above 8.7, using TruSeq Stranded mRNA library prep (Illumina, San Diego, CA, USA) according to manufacturer’s protocol. Libraries were sequenced as single-read, 50 bp on Illumina HiSeq 4000 System, 16 samples/lane, resulting in approximately 25 million reads/sample. The hippocampus (N = 95) and striatum (N =88) samples were used for gene expression-trait correlations described below. Gene expression in these brains is presumed to reflect both genetic effects and changes induced by prior ethanol treatment.

### RNA extraction and sequencing for expression QTL mapping

Brains from a separate cohort of alcohol-naïve DO mice of both sexes (M = 186, F = 183) drawn from generations G21, G22, and G23 were used for eQTL mapping. Mice were purchased from the Jackson Laboratory and transferred at wean to an intermediate barrier specific pathogen free room within Jackson Laboratory. All procedures were approved by Jackson Laboratory IACUC in accordance with NIH guidelines for the care and use of laboratory animals. Mice were individually housed under (12:12) light-dark cycle and allowed *ad libitum* access to the standard rodent chow and acidified water (pH 2.5–3.0) supplemented with vitamin K, per colony-wide standards. Mice were in cages with pine-shaving bedding (Hancock Lumber) and environmental enrichment consisting of a nestlet (Ancare, Bellmore, NY, USA) and Shepherd Shack (Shepherd Specialty Papers, Watertown, TN, USA). At 3-6 months of age, mice were phenotyped four separate times, Monday-Thursday (openfield, light-dark, holeboard, and novelty place preference) and euthanized via decapitation on Friday. Behavioral testing and euthanasia were consistently performed between 8AM-12PM. Whole intact brains were removed, hemisected and incubated in RNALater for 8-14 minutes. The striatum was removed and soaked in RNALater for a further 24 hours at room temperature and thereafter stored at −80°C. Striatum was homogenized and total RNA isolated by a TRIzol Plus kit (Life Technologies, Carlsbad, CA, USA) according to manufacturer’s method including an on-the-column DNase digestion. Quality of isolated RNA was assessed using an Agilent 2100 Bioanalyzer instrument (Agilent Technologies, Santa Clara, CA, USA) and RNA 6000 Nano LabChip assay. Ribo-depleted RNA was prepared using the RNA RiboErase Stranded cDNA library (Kapa Biosystems, Wilmington, MA, USA). Samples were pair-end sequenced on an Illumina HiSeq4000, producing paired 100 bp reads at a depth of 50 M read pairs per sample. Gene expression in these brains is presumed to reflect genetic effects independent of ethanol treatment.

### Quantitative trait locus mapping

Genome imputation, reconstruction, and QTL mapping were carried out using DOQTL and Rqtl2 software as described previously (Svenson et al. 2012; Gatti et al. 2014; Broman et al. 2019). Briefly, DOQTL and Rqtl2 software are R packages specialized for QTL mapping in multi-parent populations derived from more than two founder strains. They allow users to perform genome scans using a linear mixed model to account for population structure and permit imputation of SNPs based on founder strain genomes. Lab and generation were included as covariates for association and linkage mapping.

### Association mapping

For genome-wide association mapping, we imputed all high-quality SNPs from the Sanger Mouse Genome Project (build REL 1505; (Keane et al. 2011)) onto DO genomes and fit an additive genetic model at each SNP. This approach is widely used in human GWAS and increases power and precision by measuring the effects at individual variants by mapping at the two-state SNP level (Gatti et al. 2014). We accounted for genetic relatedness between mice by using a kinship matrix based on the leave-one-chromosome-out (LOCO) method (Cheng et al. 2013). The LOCO method was chosen because kinship calculations that include the causative marker are known to produce overly conservative mapping results (Cheng et al. 2013). Traits were rankZ transformed prior to association mapping. Genome-wide significance thresholds were calculated using 1000 permutations to create a distribution for the null hypothesis towards determining statistical significance. QTLs that exceeded *p* = 7.3 x 10^-7^ (-log_10_ *p* value = 6.14) were considered to be significant at *p* < 0.05, and QTLs that reached *p* = 1.4 x 10^-6^ (-log_10_ *p* value = 5.84) were considered to be suggestive at *p* < 0.1.

### Linkage mapping

We used an additive haplotype model with kinship correction to estimate founder effects for each QTL. Following QTL identification, we applied local SNP association to identify key SNPs in specific founder strains correlating to significant QTL effects. Traits that were not normally distributed were rankZ transformed prior to linkage mapping. The genome-wide significance thresholds for QTL were calculated using 1000 permutations to create a null distribution of LOD scores. Significant QTLs were determined at a genome-wide *p* value < 0.05 and suggestive QTLs were determined at a genome-wide *p* value < 0.1. A 95% Bayesian credible interval was used to determine the QTL region (Gatti et al. 2014; Broman et al. 2019). If a given QTL had multiple peaks (as was the case for the QTL on chromosome 2), a 1.5-LOD support interval was used to determine the QTL region instead.

### Expression-Trait correlations

Gene expression-behavior correlations were performed using Spearman’s correlations following rank normal transformations of both the gene expression and behavior; separately for each tissue type (Spearman correlations, N = 88 for striatum (∼19k genes), N = 95 for hippocampus (∼19.1k genes)). Correlations were multiple testing adjusted using the FDR approach. Expression-trait correlations with FDR < 0.25 were used in further downstream analysis.

### Expression QTL mapping

EQTL mapping was performed as described previously (Philip et al *in preparation*) using the mm10 genome assembly and Ensembl v90. Briefly, gene expression counts were obtained by summing expected counts over all transcripts for a given gene. eQTL mapping was performed on regression residuals of 17,248 genes in R/qtl2 using the founder haplotype regression method. Kinship matrices to correct for population structure were computed with the LOCO method for kinship correction (Gatti et al 2014; http://kbroman.org/qtl2). Sex and generation were included as additive covariates. We then used the interactive, web-based analysis tool QTLViewer (Philip et al *in preparation*) to visualize the expression data with profile, correlation, LOD, effect, mediation and SNP association plots. Detailed information about the structure of the QTLViewer objects are available here https://github.com/churchill-lab/qtl-viewer/blob/master/docs/QTLViewerDataStructures.md. We then determined which genes with eQTLs contained SNPs in linkage disequilibrium with the peak marker of the behavioral QTL as measured by *D*′ > .8 using the R/genetics (v1.3.8.1.3) package (*D* ′ = 0 indicative of linkage equilibrium between the two SNPs and *D* ′ = 1 representing the opposite). Finally, we tested for patterns of association between positional candidate genes and eQTLs by calculating Pearson’s correlation coefficients between the eight state founder coefficients between each behavior’s peak QTL with that of suggestive (*p* < 0.10) or significant (*p* <0 .05) *cis* eQTL for each candidate gene residing within the 1.5-LOD drop interval behavioral QTL. Multiple testing adjustments were performed using the FDR approach. EQTL-behavior QTL correlations with FDR < 0.25 were used in further downstream analysis.

### Bioinformatics analysis and candidate gene prioritization

To prioritize among candidate genes within our QTL regions, we gathered functional, phenotypic, and expression information from a variety of bioinformatics databases. First, we used the Mouse Genome Informatics (MGI; http://www.informatics.jax.org/genes.shtml) database to search for the all protein-coding genes located in our QTL support intervals and to search for genes that were associated with relevant phenotypes. We queried the NHGRI-EBI GWAS catalog to identify human homologs of the genes within our support intervals, and to determine whether they had been associated with alcohol-related phenotypes in humans. Lastly, we utilized GeneWeaver (https://www.geneweaver.org/) to identify candidate genes using a computationally predictive approach based on convergent evidence for eQTL variants and gene products associated with relevant traits. The GeneWeaver gene-sets were generated from our genomic datasets consisting of positional candidate genes (GS398277, GS398278, GS398279), genes with significant cis eQTLs (GS357751, GS357756, GS357750), and gene expression-trait correlation data from hippocampus & striatum for all genes with unadjusted *p* < 0.05 (GS398269, GS398273, GS398270, GS398274, GS398272, GS398276). We used the hierarchical similarity (HiSim) graph tool for grouping functional genomic datasets based on the genes they contained. This allowed us to identify high order intersections among genes across datasets. We then integrated information gleaned across these multiple bioinformatics platforms with our QTL, eQTL, and gene expression-trait correlation data to generate a list of prioritized genes based upon multiple lines of supporting evidence.

## Results

### Phenotypic analysis

We observed large amounts of variation in ethanol-induced ataxia, ethanol-induced hypothermia, and ethanol-induced LORR (**Supplementary Figures 1, 2, & 3)**. In addition, we observed small but significant correlations among the three traits (**Supplementary Figure 4**). Although the effect sizes of the correlations were quite small, our results suggest that these three traits do not have completely distinct neurogenetic underpinnings. Additional phenotypic analyses are provided in the supplementary materials and figures.

### QTL analysis

We performed a QTL analysis for three measures of ethanol sensitivity (ataxia, hypothermia, LORR). We identified a significant QTL on chromosome 16 for ethanol-induced ataxia (Figure 1) and a significant QTL on chromosome 1 for ethanol-induced hypothermia (Figure 2). In addition, we identified a suggestive QTL on chromosome 2 for ethanol-induced LORR (Figure 3; both QTL analyses for LORR -- all mice vs only mice with LORR durations > 1 minute -- identified a suggestive QTL at the same location on chromosome 2). Table 1 displays the confidence interval, width, peak position, LOD scores, number of protein coding genes in the interval, and percent variance explained for each QTL. The width of the QTLs ranged between 4.53 and 6.87 Mb. Figures 4, 5, **and** 6 display the LOD scores of SNPs within each of the identified QTLs, as well as the SNP marker associations across the eight DO founders. We also estimated founder haplotype effects for each QTL. For the ethanol-induced ataxia QTL on chromosome 16, the CAST/EiJ founder haplotype was associated with enhanced ataxia following ethanol administration (Figure 7), whereas 129S1/SvlmJ, NZO/HiLtJ, and WSB/EiJ haplotypes were associated with decreased ataxia following ethanol administration, and A/J, NOD/ShiLtJ, PWK/PhJ, C57BL/6J haplotypes fell in between. Together, the haplotype patterns (Figure 7) and SNP marker associations (Figure 4) across the eight DO founder strains suggest that there may be at least three distinct haplotypes underlying the chromosome 16 QTL. For the ethanol-induced hypothermia QTL on chromosome 1, the C57BL/6J founder haplotype was associated with decreased body temperatures post-ethanol (Figure 8). This pattern was also observed in the SNP marker associations of the eight DO founders (Figure 5), thus suggesting that DO mice with C57BL/6J alleles at this locus were more sensitive to the hypothermic effects of ethanol. Finally, for the ethanol-induced LORR QTL on chromosome 2, the CAST/EiJ founder haplotype (Figure 9) and SNPs (Figure 6) were associated with decreased sensitivity to the sedative effects of ethanol as demonstrated by decreased LORR duration.

**Figure 1.**
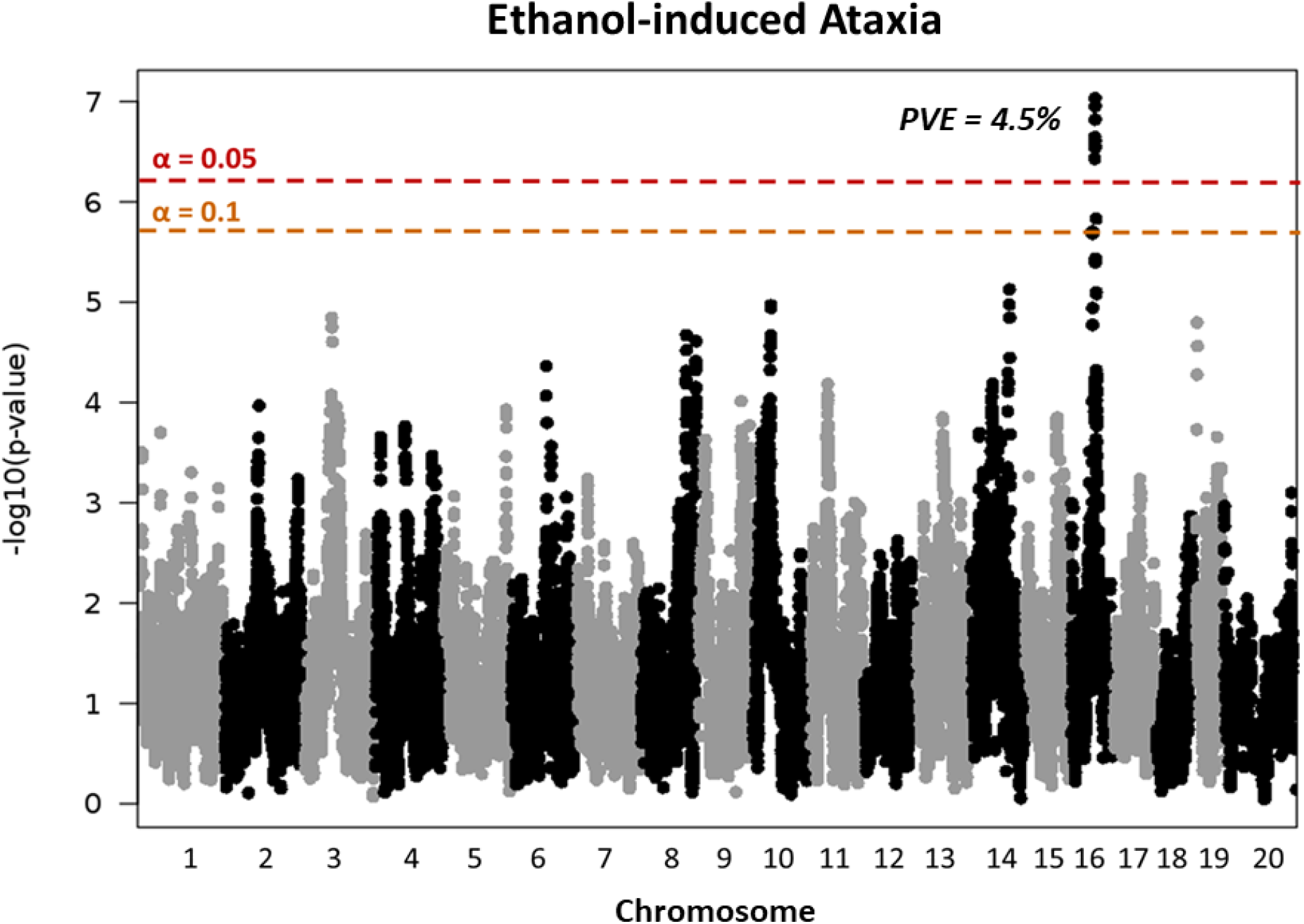
Genome-wide association mapping of ethanol-induced ataxia in DO mice. There was a significant QTL on chromosome 16 for ethanol-induced ataxia. Each point plots the –log10 *p*-value for the association between one SNP and ataxia index score. Values were plotted against their respective positions on the chromosomes. The dashed red and orange lines represent the genome-wide significant (*p* < 0.05) and genome-wide suggestive (*p* < 0.1) thresholds, respectively. PVE = percent variance explained.

**Figure 2.**
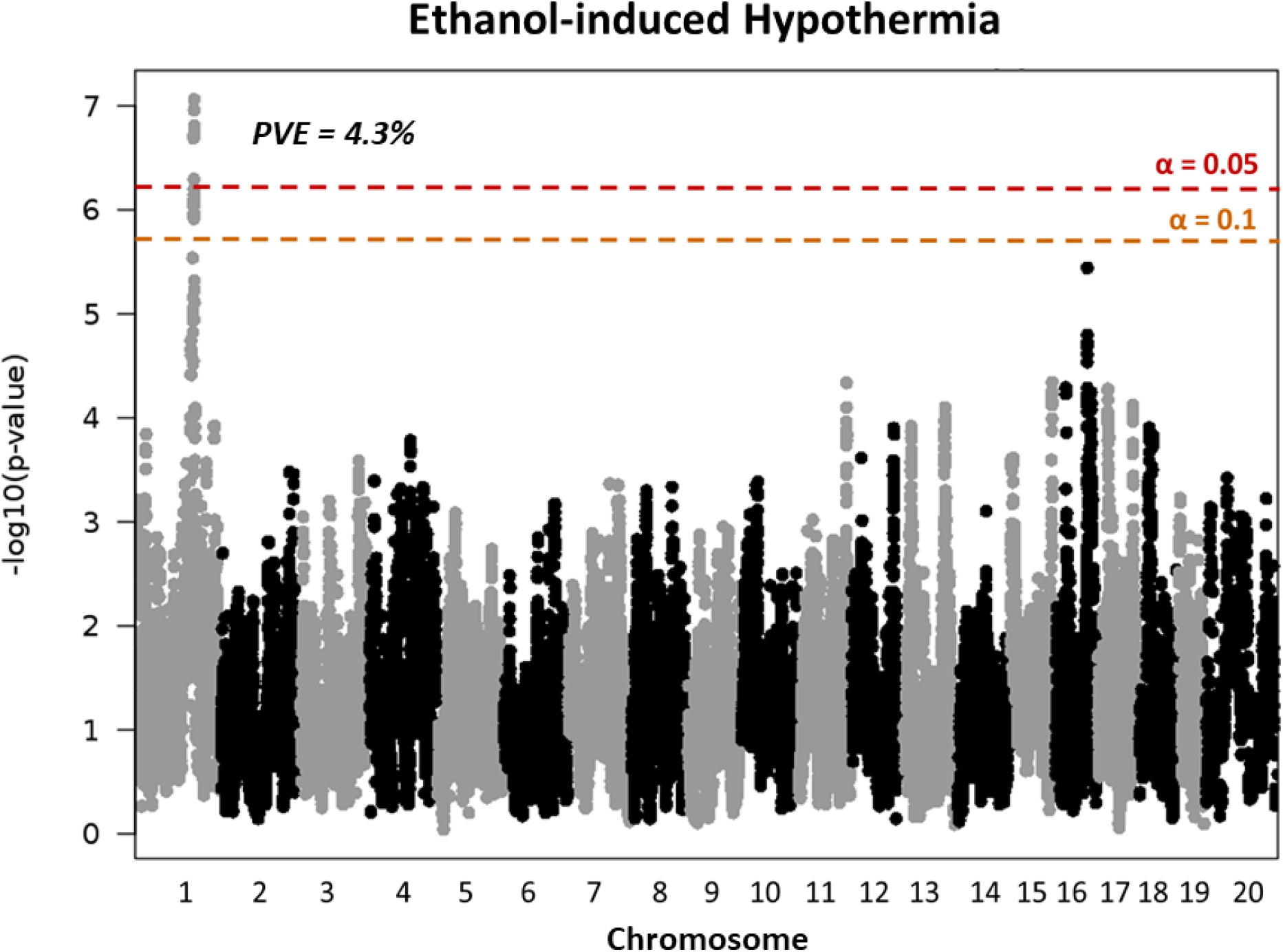
Genome-wide association mapping of ethanol-induced hypothermia in DO mice. There was a significant QTL on chromosome 1 for ethanol-induced hypothermia. Each point plots the –log10 *p*-value for the association between one SNP and hypothermia index score. Values were plotted against their respective positions on the chromosomes. The dashed red and orange lines represent the genome-wide significant (p < 0.05) and genome-wide suggestive (p < 0.1) thresholds, respectively. PVE = percent variance explained.

**Figure 3.**
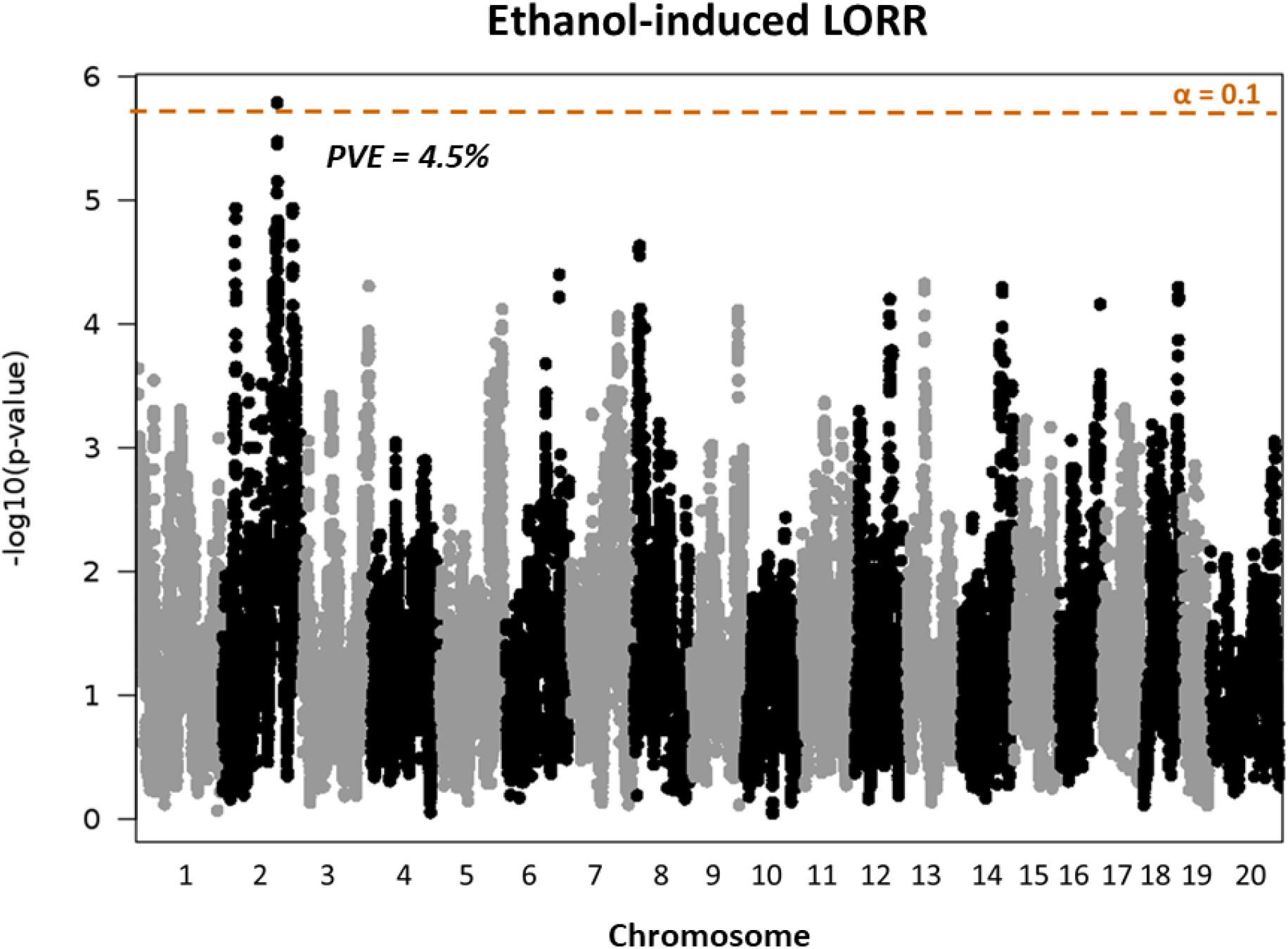
Genome-wide association mapping of ethanol-induced LORR in DO mice. There was a suggestive QTL on chromosome 2 for ethanol-induce LORR. Each point plots the –log10 *p-*value for the association between one SNP and LORR index score. Values were plotted against their respective positions on the chromosomes. The dashed orange line represents the genome-wide suggestive (p < 0.1) threshold. PVE = percent variance explained.

**Figure 4.**
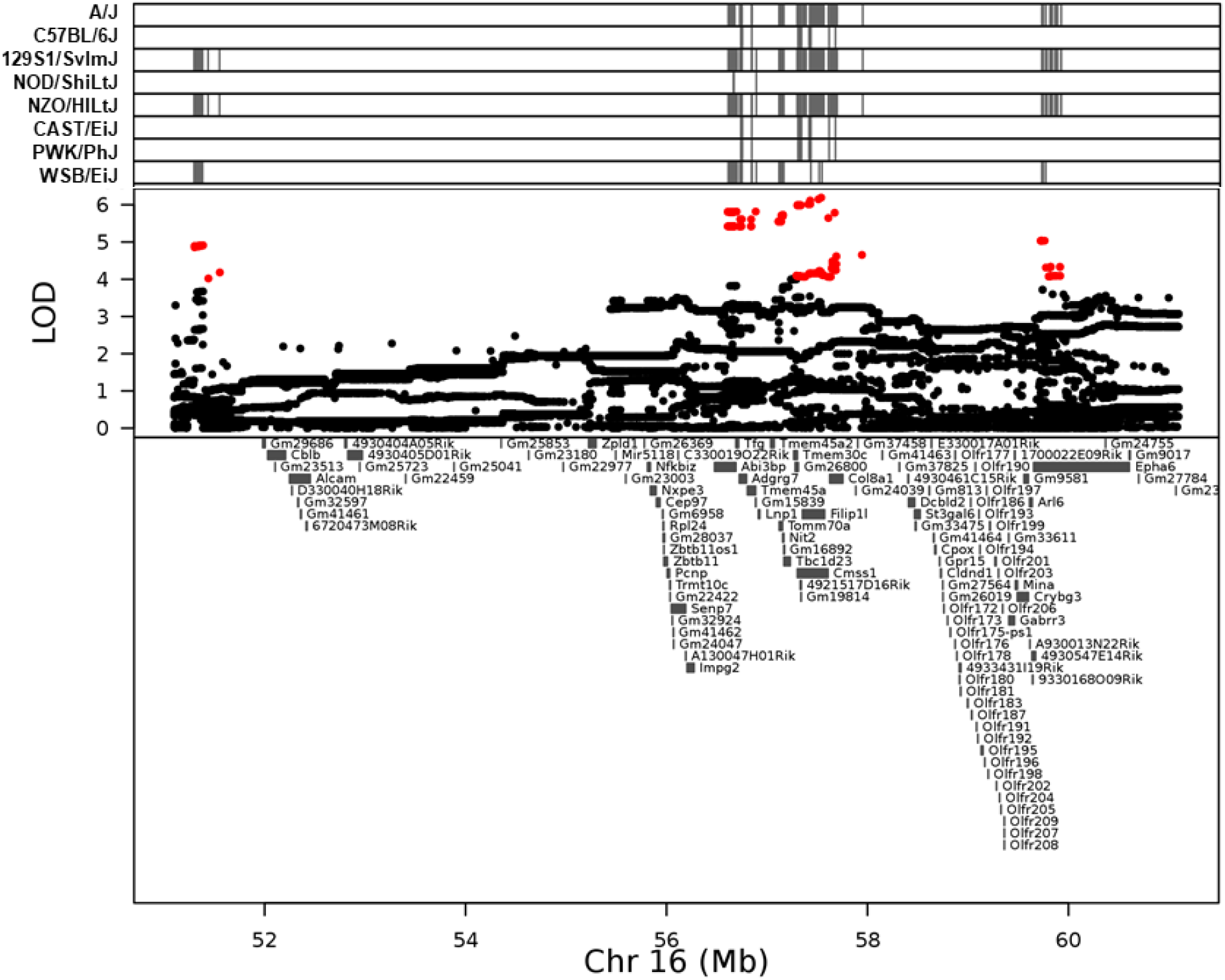
Association mapping of QTL for ethanol-induced ataxia in DO mice. Top panel shows SNP marker associations of the eight DO founders. The x-axis displays the distribution along the chromosome in physical distance. The y-axis displays the LOD score for the SNP mapping. Each data point shows the LOD value at one SNP, red data points indicate scores above the *p* < 0.1 threshold. Below each plot is an expanded view showing known genes within the QTL support interval (53.60582 – 60.47718 Mb).

**Figure 5.**
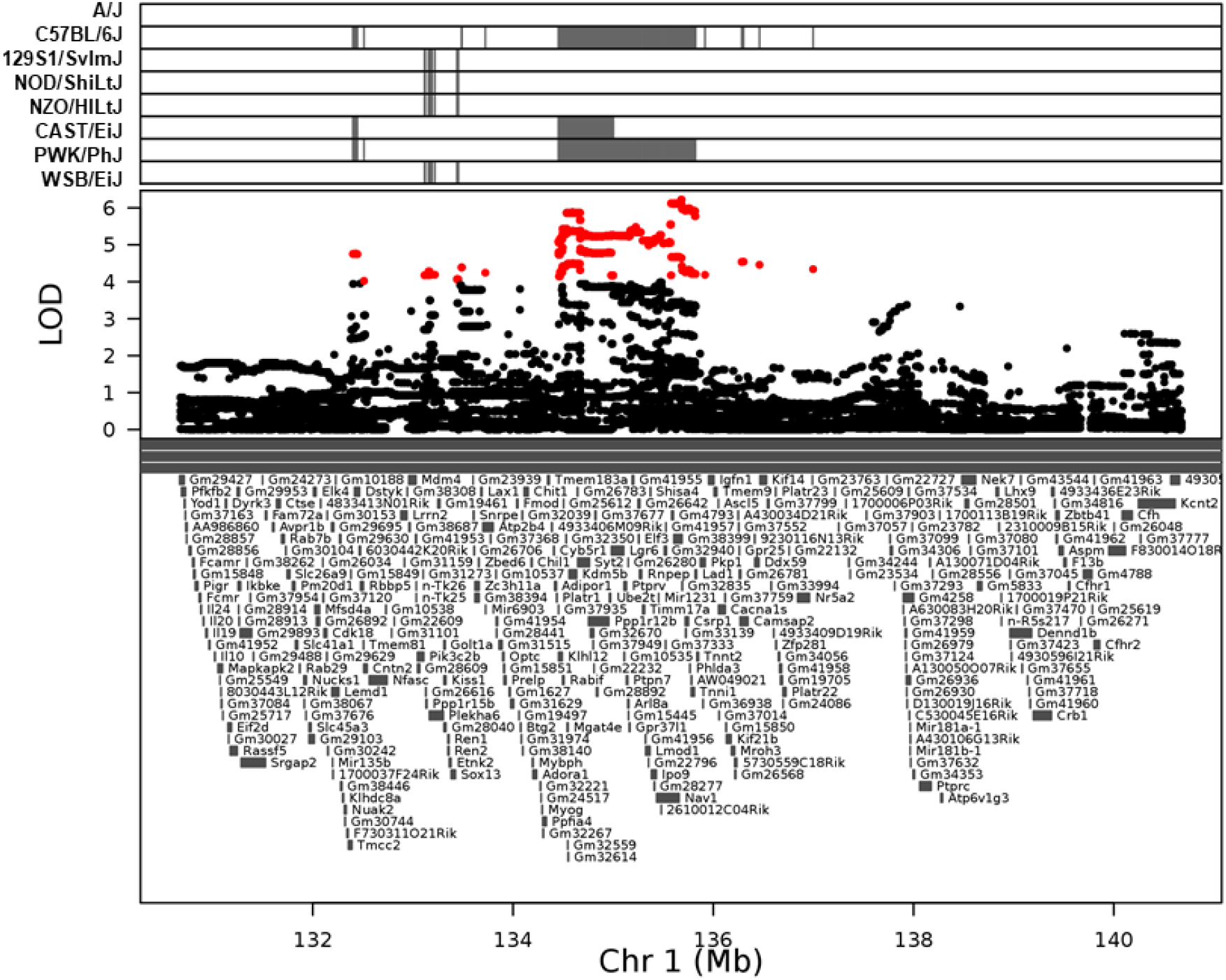
Association mapping of QTL for ethanol-induced hypothermia in DO mice. Top panel shows SNP marker associations of the eight DO founders. The x-axis displays the distribution along the chromosome in physical distance. The y-axis displays the LOD score for the SNP mapping. Each data point shows the LOD value at one SNP, red data points indicate scores above the *p* < 0.1 threshold. Below each plot is an expanded view showing known genes within the QTL support interval (132.6132 – 137.1381).

**Figure 6.**
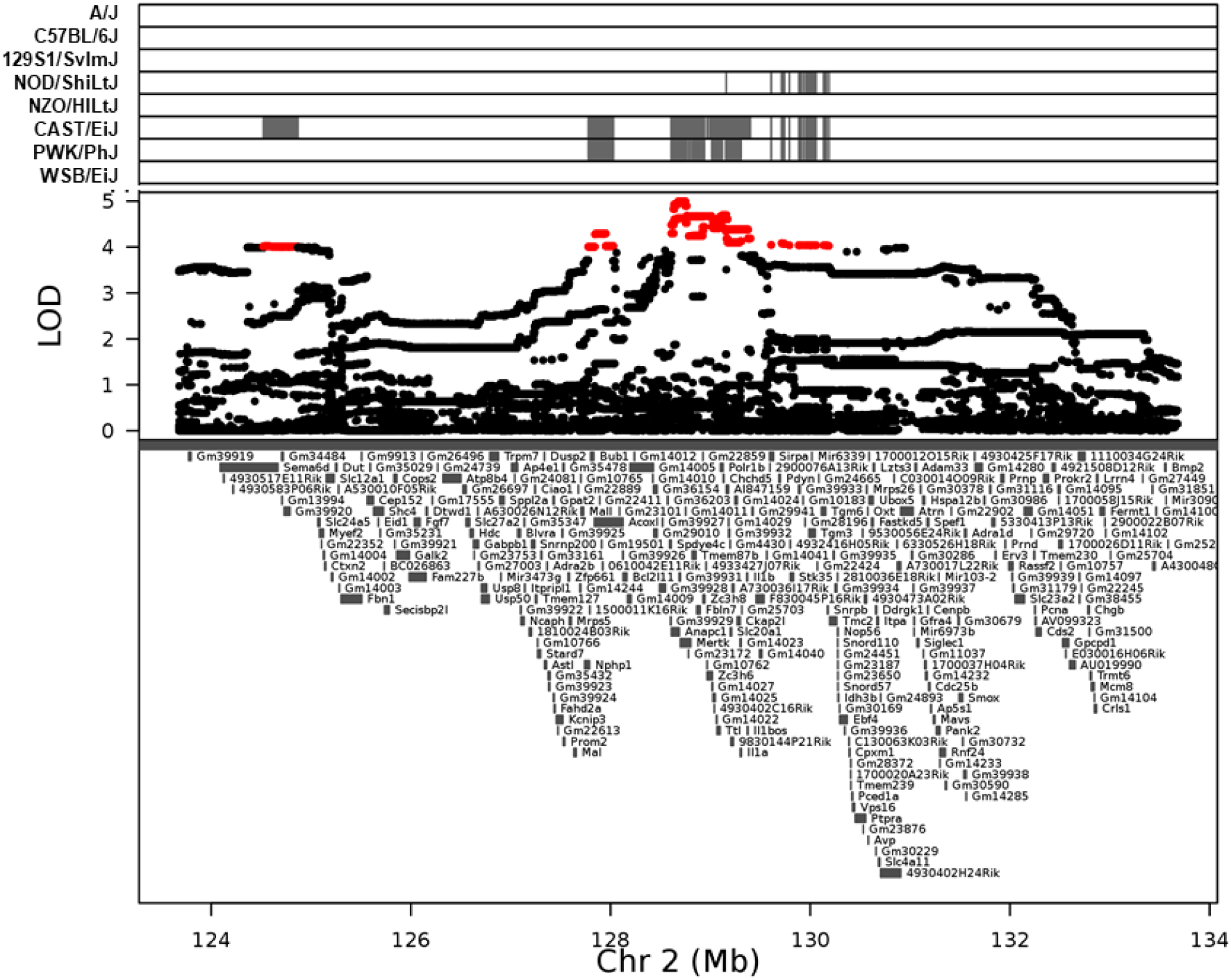
Association mapping of QTL for ethanol-induced LORR in DO mice. Top panel shows SNP marker associations of the eight DO founders. The x-axis displays the distribution along the chromosome in physical distance. The y-axis displays the LOD score for the SNP mapping. Each data point shows the LOD value at one SNP, red data points indicate scores above the *p* < 0.1 threshold. Below each plot is an expanded view showing known genes within the QTL support interval (124.9770 – 129.7152 Mb).

**Figure 7.**
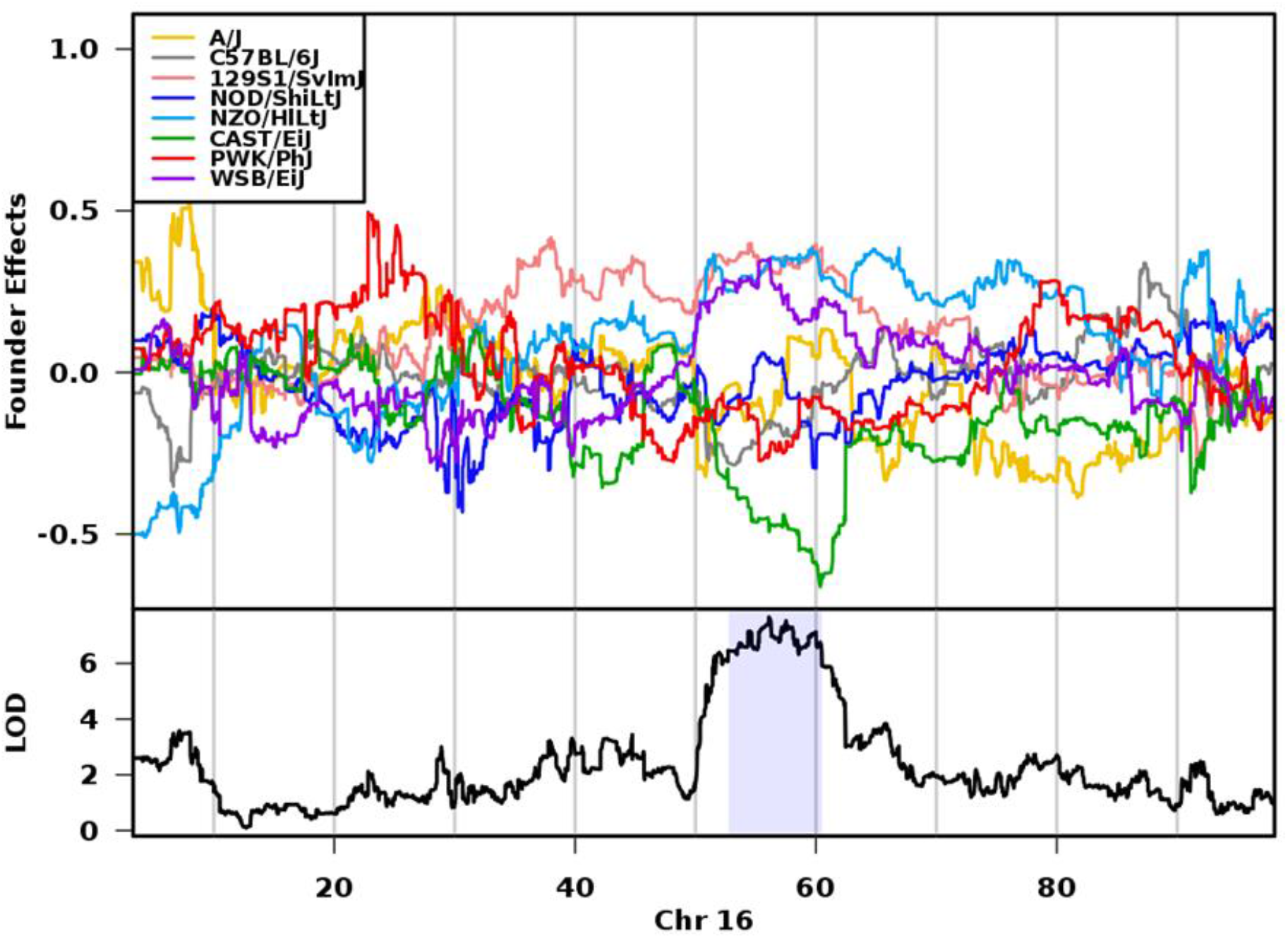
Haplotype effects on chromosome 16 QTL for ethanol-induced ataxia in DO mice. The top panel shows the eight DO founder allele effects determined by linkage mapping. The x-axis is physical distance in Mb along the chromosome. The y-axis for the top panel is the effect coefficient, and the bottom panel is the LOD score. At ∼56 Mb, the CAST/EiJ (green) alleles are associated with enhanced ataxia following ethanol administration, the 129S1/SvlmJ (pink), NZO/HiLtJ (light blue), and WSB/EiJ (purple) alleles are associated with decreased ataxia following ethanol administration, and A/J (yellow), NOD/ShiLtJ (dark blue), PWK/PhJ (red), and C57BL/6J (grey) alleles were in between. Shading in the bottom panel identifies the 95% credible interval.

**Figure 8.**
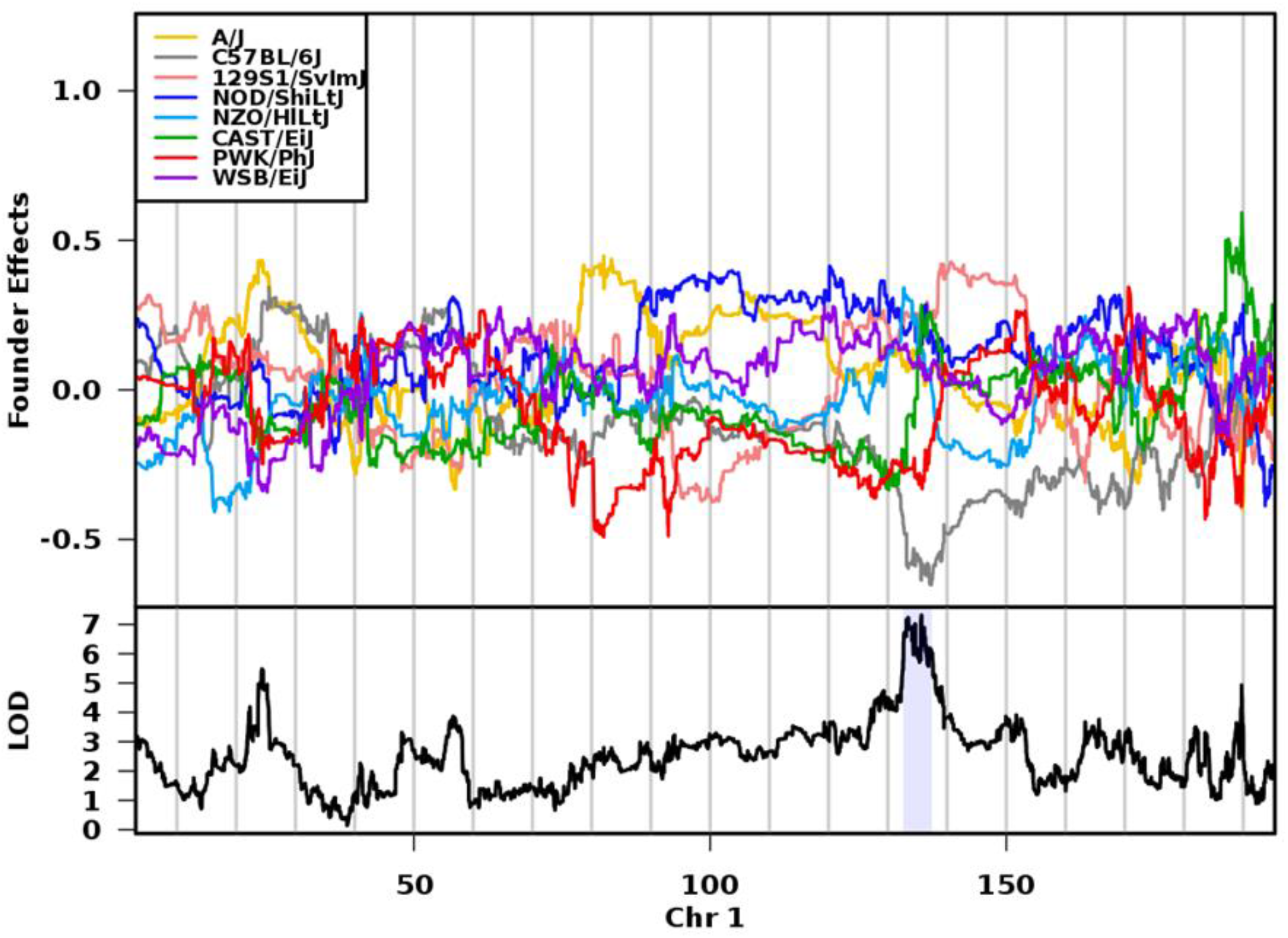
Haplotype effects on chromosome 1 QTL for ethanol-induced hypothermia in DO mice. The top panel shows the eight DO founder allele effects determined by linkage mapping. The x-axis is physical distance in Mb along the chromosome. The y-axis for the top panel is the effect coefficient, and the bottom panel is the LOD score. At ∼135 Mb, the C57BL/6J alleles (grey) are associated with decreased body temperature following ethanol administration. Shading in the bottom panel identifies the 95% credible interval.

**Figure 9.**
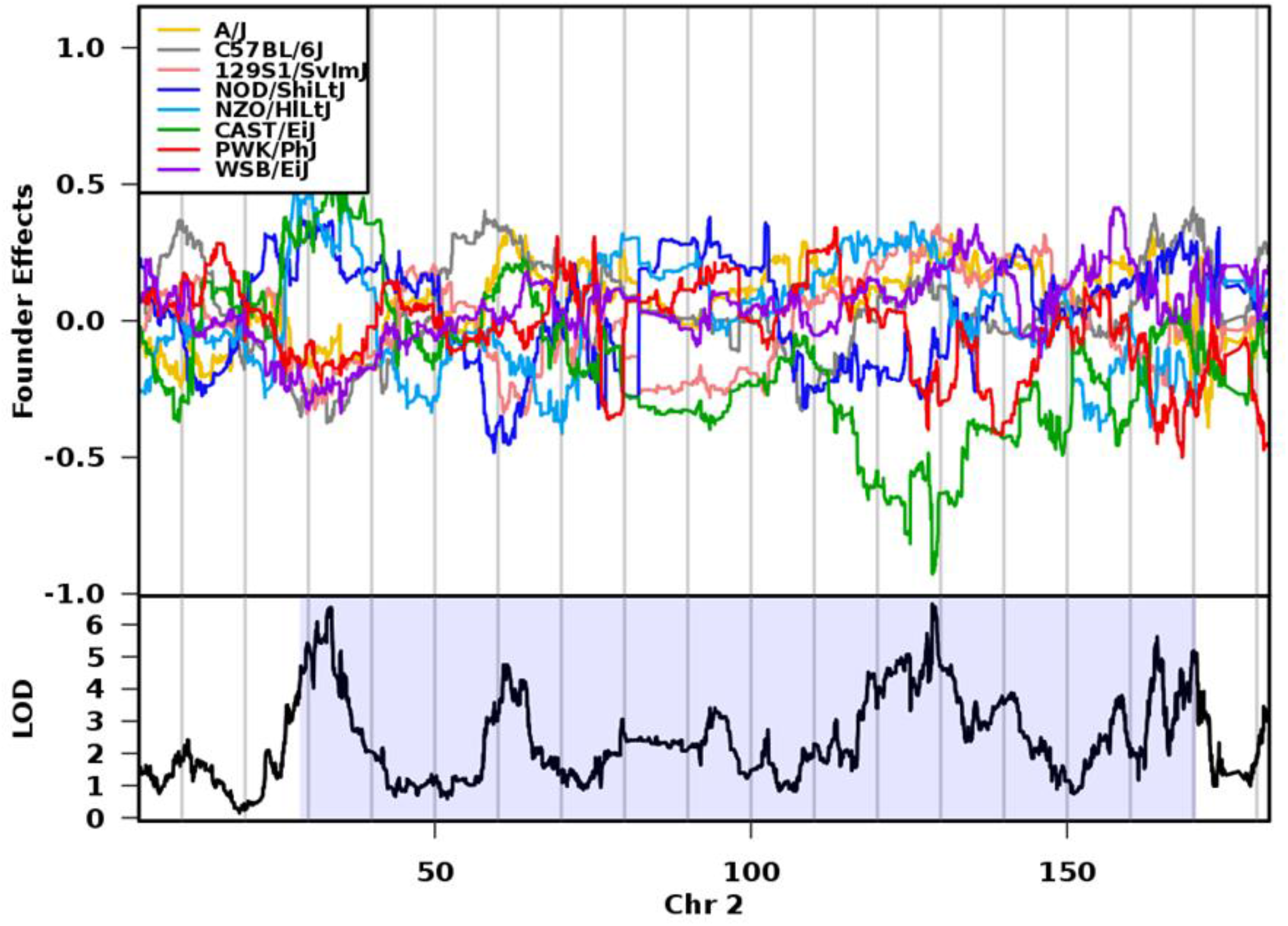
Haplotype effects on chromosome 2 QTL for ethanol-induced LORR in DO mice. The top panel shows the eight DO founder allele effects determined by linkage mapping. The x-axis is physical distance in Mb along the chromosome. The y-axis for the top panel is the effect coefficient, and the bottom panel is the LOD score. At ∼129 Mb, the CAST/EiJ alleles (green) are associated with decreased duration of LORR following ethanol administration. Shading in the bottom panel identifies the 95% credible interval.

**Table 1.** Ethanol sensitivity QTLs. The confidence interval (CI and peak marker location are given in Mb position based on Build 38 of the mouse chromosome.

| QTL | Chr | CI (Mb) | Width (Mb) | Peak (Mb) | LOD score | Protein coding genes in interval | % variance explained |
| --- | --- | --- | --- | --- | --- | --- | --- |
| **Ataxia** |  |  |  |  |  |  |  |
| *Difference in latency to fall between pre- and post- treatment* | 16 | 53.60582 – 60.47718 | 6.87 | 56.10037 | 8.38 | 61 | 4.5% |
| **Hypothermia** |  |  |  |  |  |  |  |
| *Average temperature post-ethanol* | 1 | 132.6132 – 137.1381 | 4.53 | 135.6791 | 7.91 | 72 | 4.3% |
| **Loss of Righting Response** |  |  |  |  |  |  |  |
| *Duration of LORR with mice with zero sleeptime excluded* | 2 | 124.9770 – 129.7152 | 4.73 | 128.7657 | 7.31 | 69 | 4.5% |

### Expression-trait correlations

Next we searched for correlations between striatal and hippocampal gene expression levels for genes located within our QTLs and ethanol traits in the DO mice (**Tables S1 & S2**). Using an FDR corrected threshold of *p* < 0.25, we identified three genes with striatal expression levels that were correlated with ethanol-induced ataxia (*Deup1, Pcdhga7*, and *Pcdhgb4*). However, we did not observe any additional expression-trait correlations that reached the selected FDR corrected threshold, likely due to the small number of available tissue samples (striatum N = 88, hippocampus N = 95).

### Expression QTL mapping

We performed eQTL mapping to determine which positional candidate genes had striatal expression levels that mapped to the same interval (± 2 Mb) as the behavioral QTL (Table 2). We further prioritized among these eQTL genes by: 1) determining which genes contained SNPs that were in linkage disequilibrium with the peak marker as measured by *D*′ > .8, and 2) identifying which genes displayed expression patterns that were significantly correlated (*p* < 0.05) with the haplotype effects. Within the ataxia QTL on chromosome 16, 19 genes had significant eQTLs. Ten of those genes possessed SNPs with *D*′ > .8 (*Cep97, Tbc1d23, Impg2, Nfkbiz, Abi3bp, Cldnd1, Riox2, Cpox, Senp7, Pcnp*) and three (*Cpox, R* = .72; *Arl6, R* = .74; and *Riox2, R* = .72) showed expression patterns that were significantly correlated with the founder haplotype effects. For the hypothermia QTL on chromosome 1, 26 genes had significant eQTLs (*p* < 0.05) that mapped to the same region as the behavioral QTL. Of those genes, 14 contained SNPs with *D*′ > .8 (*Phlda3, Gm37333, Lmod1, Ipo9, Ptpn7, Ppp1r12b, Nav1, Gm26781, Arl8a, Rnpep, Zbed6, Zfp281, Gm37949, Chil1*); and five in particular (*Nav1, R* = -.71; *Lmod1, R* = -.94; *Klhl12, R* = - .74; *Arl8a, R* = -.85; and *Rnpep, R* = -.73) showed haplotypic expression patterns that were significantly correlated with the haplotypic behavioral effects. For the LORR QTL, 37 genes had significant eQTLs that co-mapped with the behavioral QTL on chromosome 2. Among those, 12 genes contained SNPs with *D*′ > .8 (*Fbln7, Anapc1, Mertk, Gm10762, Zc3h8, Fahd2a, Zfp661, Pdyn, Sppl2a, Gm14212, Eid1, Cep152*), and seven of those genes (*Slc27a2, R* = .94; *Sppl2a, R* = .76; *Anapc1, R* = .73; *Mertk, R* = .75; *Fahd2a, R* = .77; *Cep152, R* = -.71; and *Mrps5, R* = .79) displayed expression patterns were significantly correlated with the founder haplotype effects.

**Table 2:**
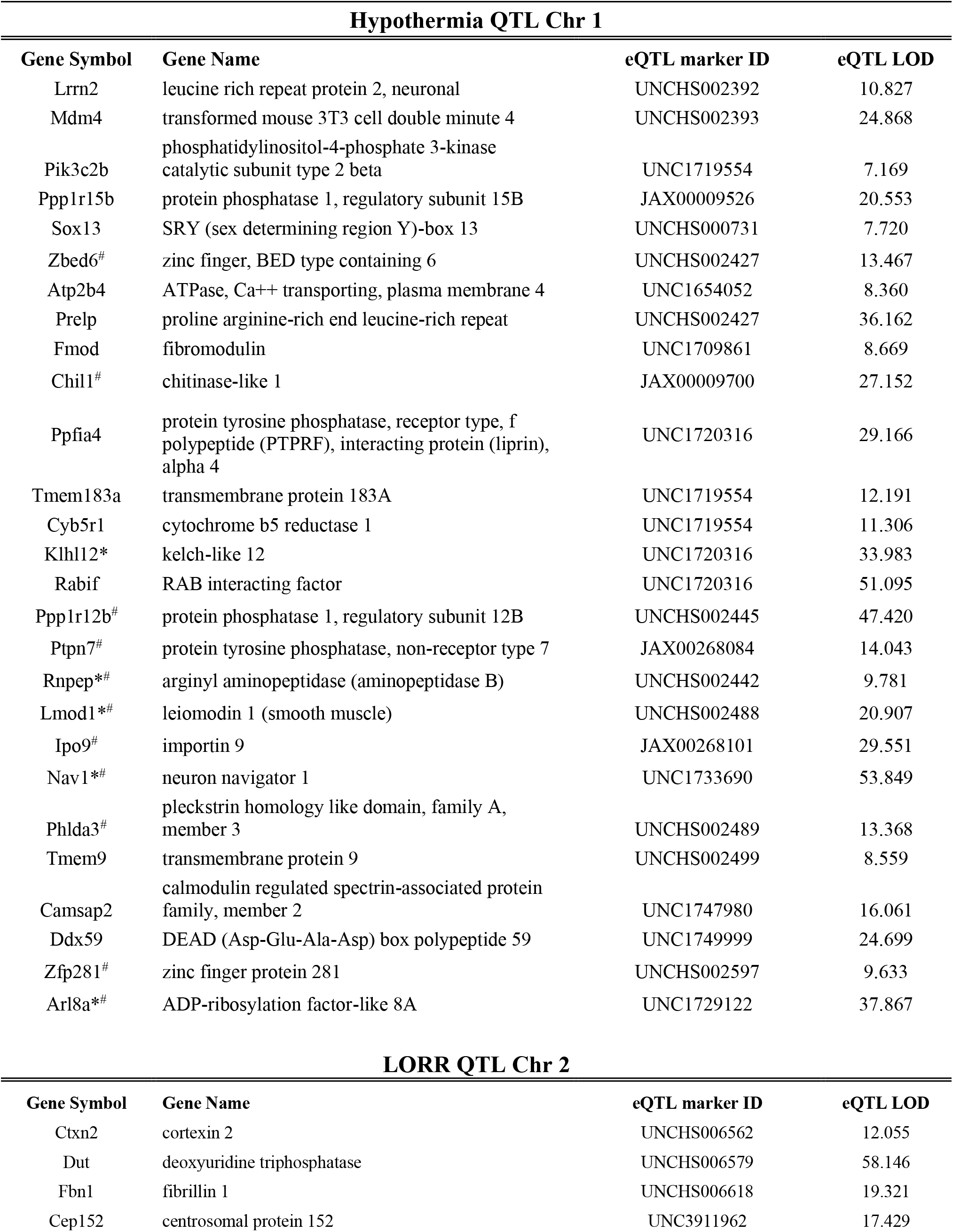
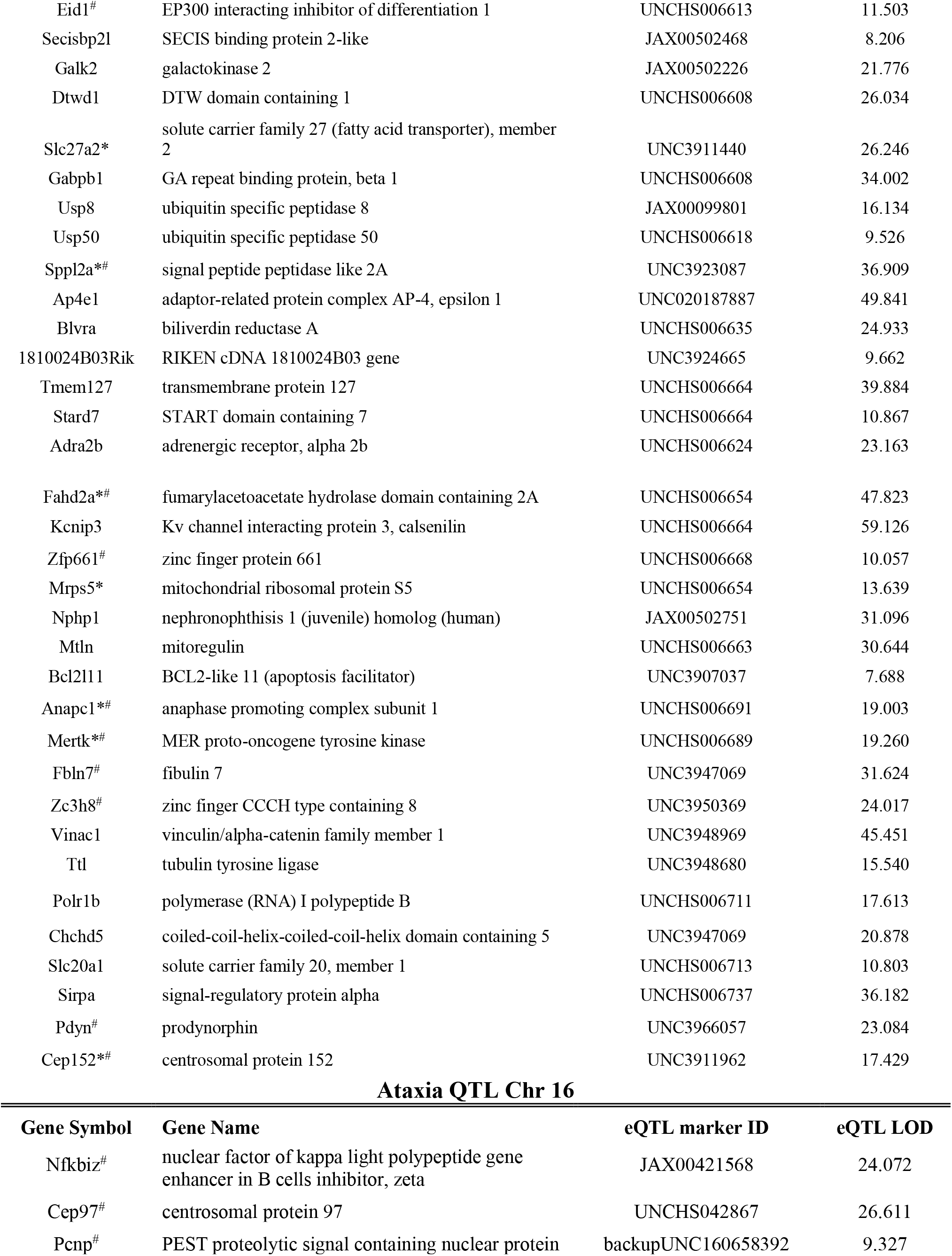
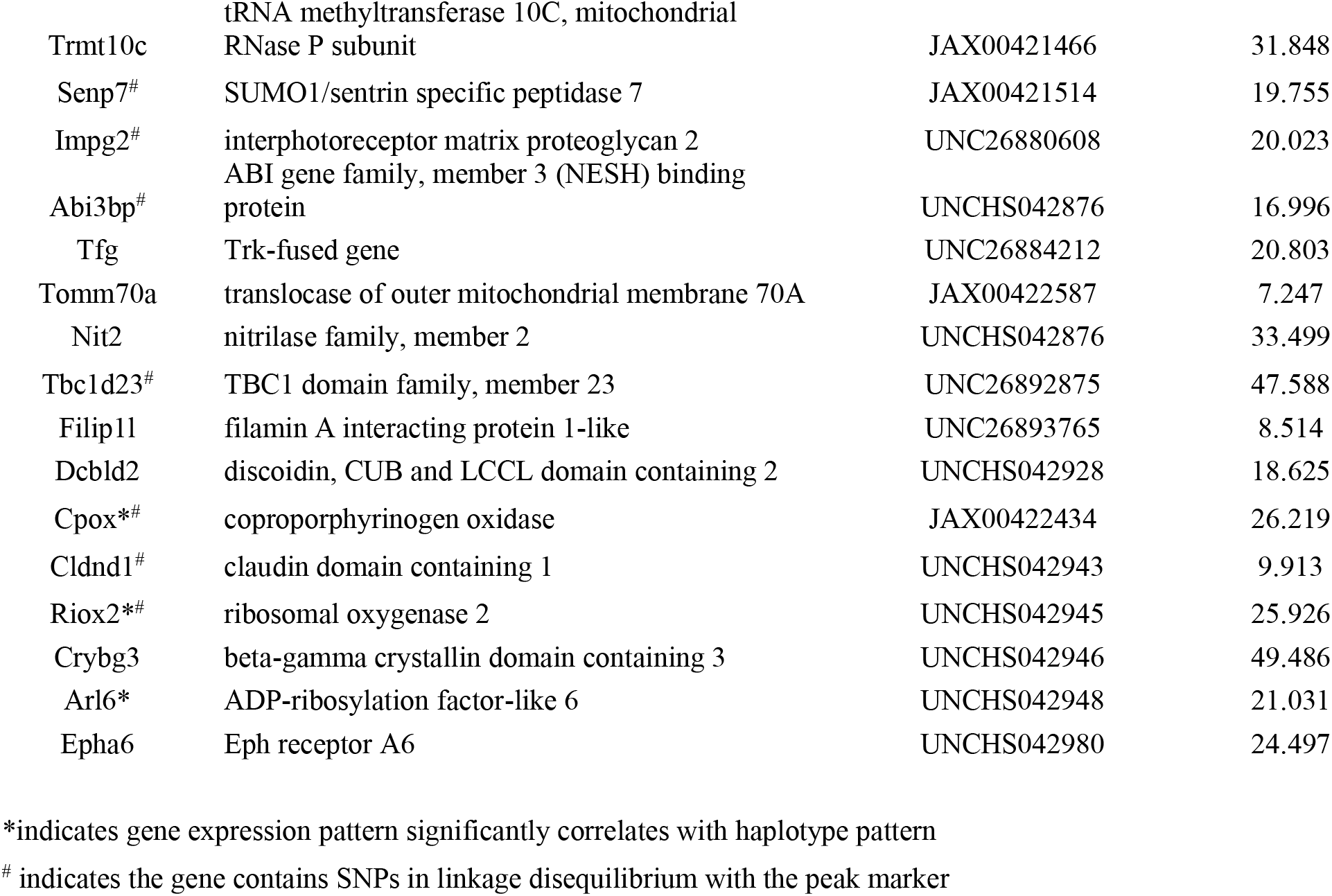
Genes with significant cis-eQTLs for ethanol-related traits in DO mice

### Bioinformatics analysis and candidate gene prioritization

We interrogated multiple independent and complementary databases to prioritize the genes within our QTL intervals (**Tables S3, S4, and S5**). Many of these genes have been associated with peripherally relevant traits of interest, some of which are trait specific (e.g., vestibular reflexes, body temperature regulation, sedation), and others are more broadly related to drug and ethanol phenotypes. We also identified a number of genes within our QTL intervals that have been previously implicated in human GWAS for drug and alcohol-related traits. Lastly, we used GeneWeaver to identify genes that were common across all of our datasets. For ethanol-induced ataxia, three genes (*Tomm70a, Pcnp,* and *Abi3bp*) were shared between gene-sets for positional candidates, cis-eQTLs, and the striatal expression-trait correlations, and one gene (*Trmt10c*) was shared between gene-sets for positional candidates, cis-eQTLs, and the hippocampal expression-trait correlations **(Supplementary Figure 8**). For ethanol-induced hypothermia, we identified one gene (*Klhl12*) that was shared between gene-sets for positional candidates, cis-eQTLs, and the hippocampal expression-trait correlations. Three genes (*Tnnt2, Adora1,* and *Gm4204*) were shared between gene-sets for chromosome 1 positional candidates and striatal expression-trait correlations, but not cis-eQTLs. Five genes (*Gm37759, 9230116N13Rik, Klhl12, Mybph*, and *Gm22232*) were shared between gene-sets for chromosome 1 positional candidates and hippocampal expression-trait correlations, but not cis-eQTLs. In addition, three genes (*Lamc2, Dpp10,* and *Creg2*) were shared between gene-sets for both hippocampal and striatal expression-trait correlations and cis-eQTLs (**Supplementary Figure 9**). For ethanol-induced LORR, we identified one gene (*Prkcq*) that was shared between gene-sets for both hippocampal and striatal expression-trait correlations and cis-eQTLs. One gene (*Bub1*) was shared between gene-sets for chromosome 2 positional candidates and striatal expression-trait correlations, and one gene (*Itpripl1*) was shared between gene-sets for chromosome 2 positional candidates and hippocampal expression-trait correlations (**Supplementary Figure 10**). We integrated all of the data described above to generate a list of the most promising candidate genes within each interval (Tables 3, 4, **and** 5).

**Table 3.** Prioritized list of candidate genes - Ataxia Chromosome 16

| Gene | cis eQTL | $D' > .8$ | Highly connected across GeneWeaver datasets | Expressed in Brain | Linked to ethanol or other relevant phenotype in rodents | Implicated in human SUD GWAS |
| --- | --- | --- | --- | --- | --- | --- |
| *Abi3bp* | Yes | Yes | Yes | Yes |  | Alcohol dependence (Mbarek et al 2015) |
| *Arl6* | **Yes** |  |  | Yes |  | Serum biomarker for excessive alcohol use (Liangpunsakul et al 2015) |
| *Cpox* | **Yes** | Yes |  | Yes | Ethanol withdrawal severity in BXD mice (Buck et al 1997) | Differential expression in frontal cortex and hippocampus in alcoholism (Liu et al 2006; Zhou et al 2011; Farris et al 2015) |
| *Pcnp* | Yes | Yes | Yes | Yes |  |  |
| *Riox2* | **Yes** | Yes |  |  |  |  |
| *Senp7* | Yes | Yes |  | Yes |  | Alcohol consumption (Linner et al 2019); Bitter alcohol consumption (Zhong et al 2019) |
| *Tbc1d23* | Yes | Yes |  |  |  |  |
| *Tomm70a* | Yes |  | Yes | Yes | Differential expression in nAcc, PFC, & VTA 4 hrs post ethanol in C57BL/6J & DBA/2J mice (Kerns et al 2005); differential expression in striatum & cortex of C57BL/6J mice drinking to intoxication (Mulligan et al 2011) |  |
| *Trmt10c* | Yes |  | Yes | Yes | Differential expression in nAcc 24 hours post ethanol access in alcohol preferring rats (Bell et al 2009) |  |
| *Zpld1* |  |  |  | Yes | Abnormal vestibuloocular reflex/system physiology, impaired swimming (Vijayakumar et al 2019) | Alcohol consumption measurement interaction with waist-hip ratio (Velez Edwards et al 2012) |
| cis-QTL indicates if the gene also had a significant cis-QTL; and those in bold had expression patterns that significantly correlated with the haplotype effects for each QTL |  |  |  |  |  |  |
| *D'* > .8 indicates a gene with an eQTL that contained SNPs in linkage disequilibrium with the peak marker |  |  |  |  |  |  |

**Table 4.**
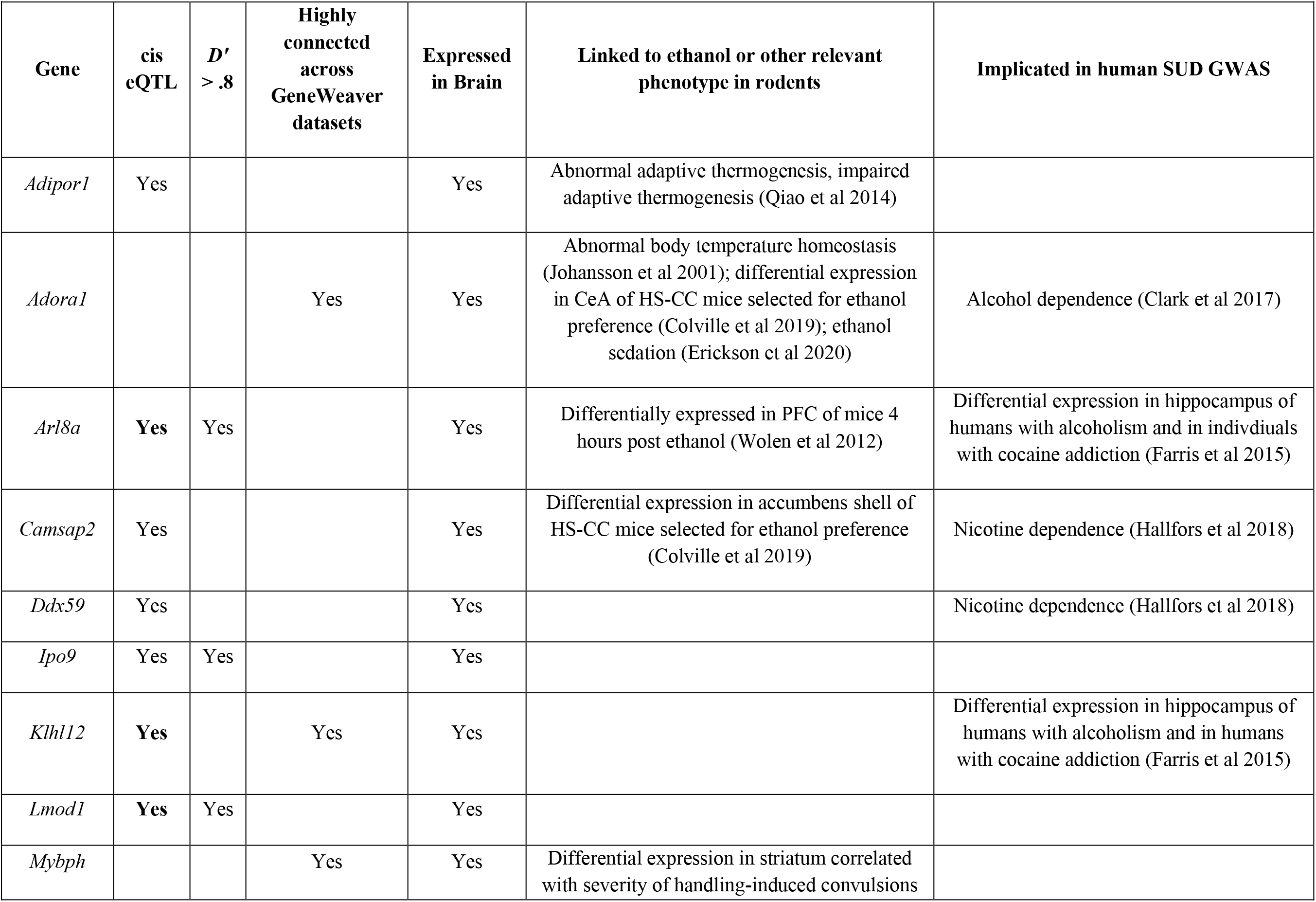
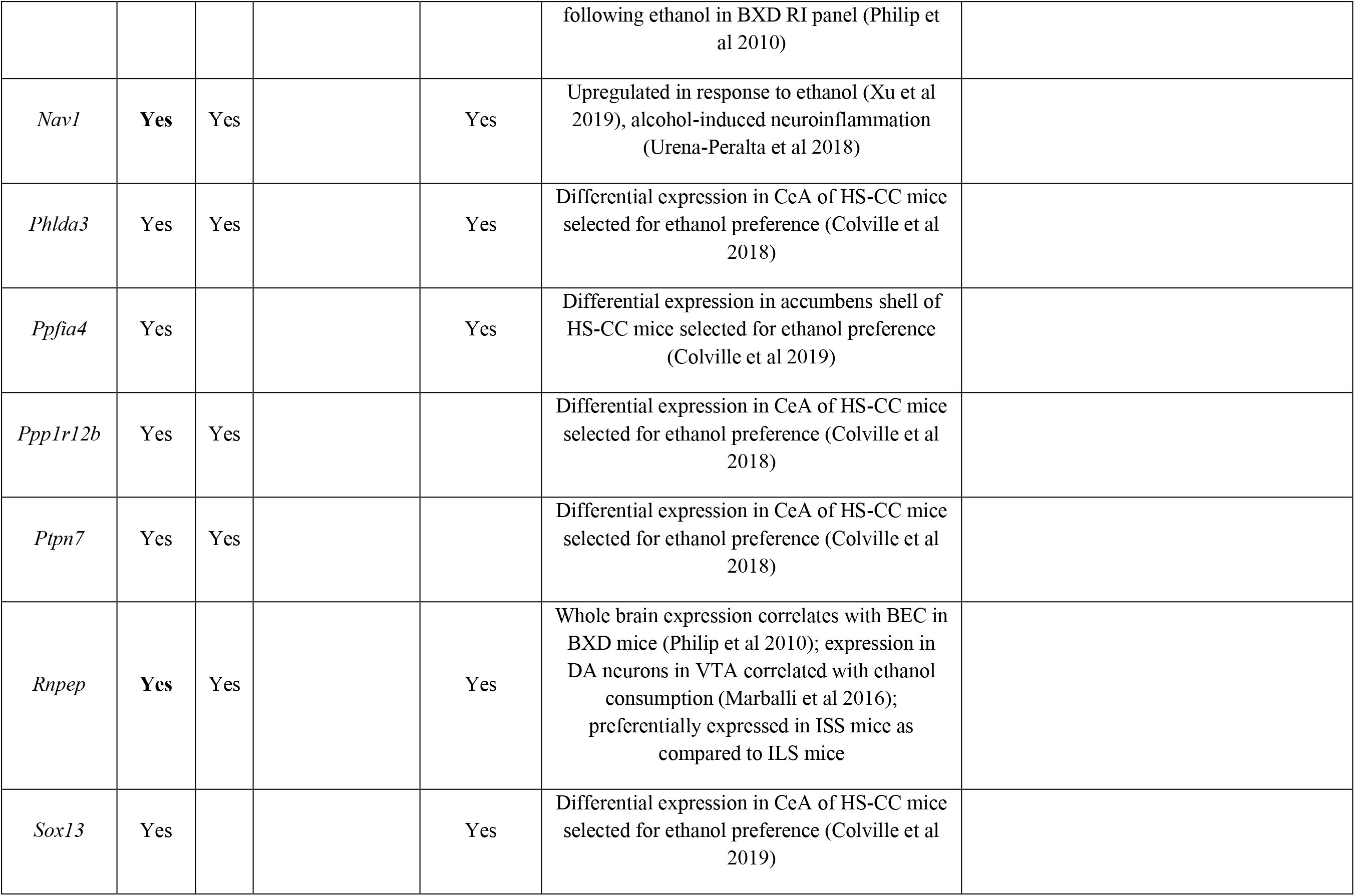
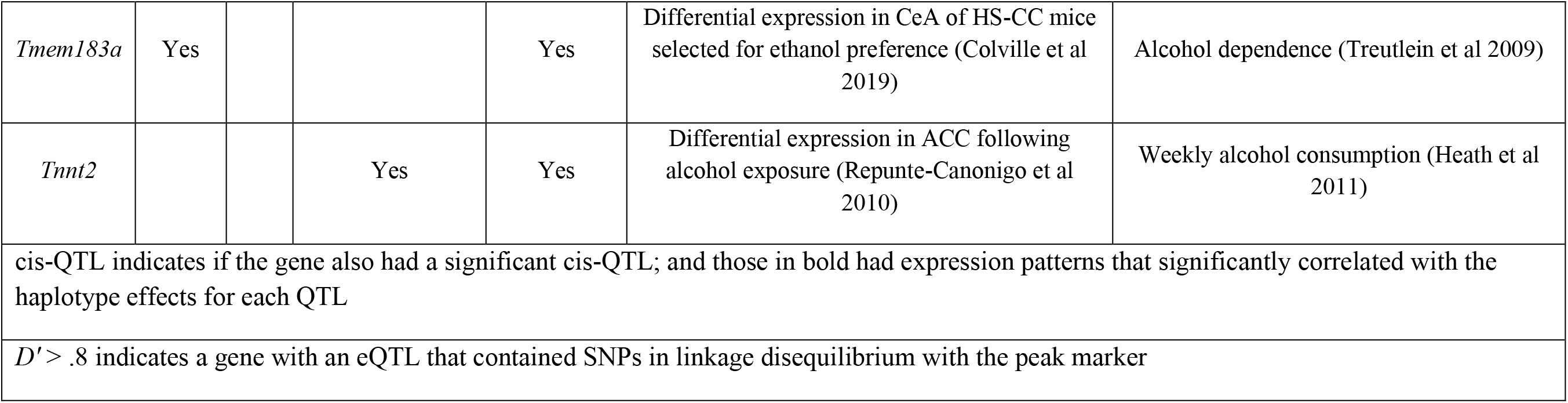
Prioritized list of candidate genes - Hypothermia Chromosome 1

**Table 5.**
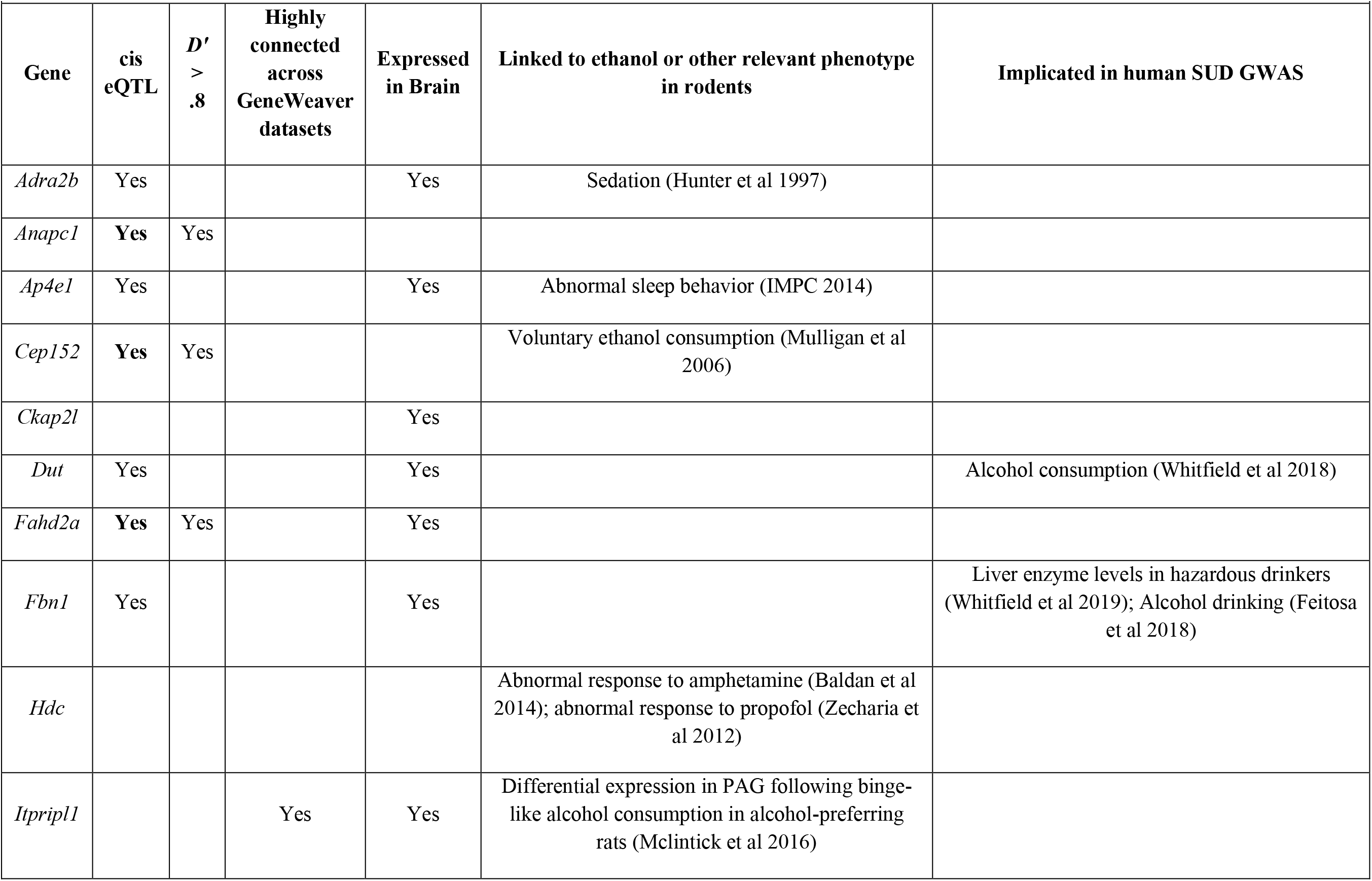
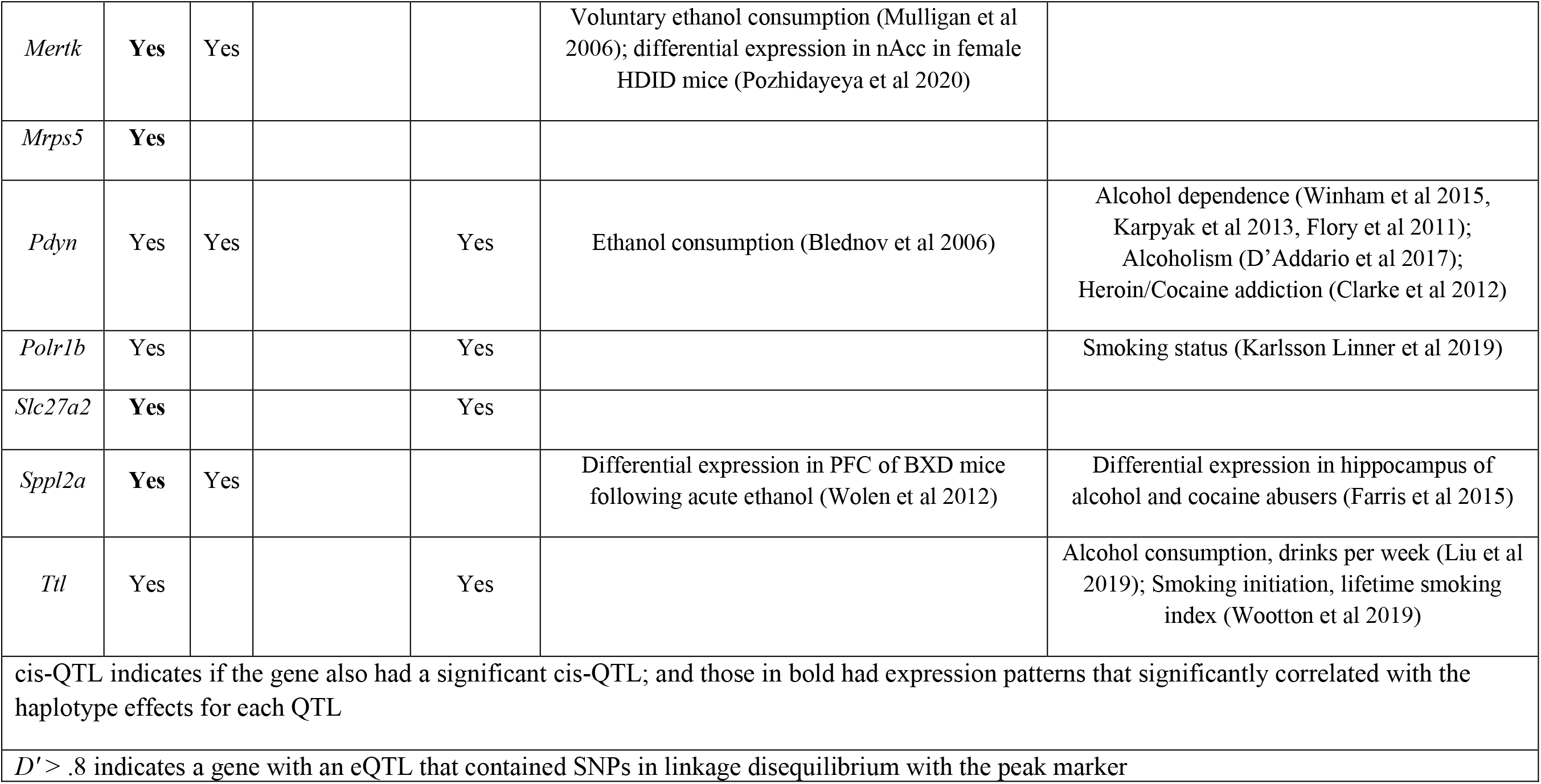
Prioritized list of candidate genes - LORR Chromosome 2

## Discussion

### QTL mapping

We performed genome-wide SNP association mapping in conjunction with linkage analysis to identify narrow genomic regions associated with ethanol sensitivity in the DO mouse population. We identified a significant QTL on chromosome 16 associated with ethanol-induced ataxia, a significant QTL on chromosome 1 associated with ethanol-induced hypothermia, and a suggestive QTL on chromosome 2 associated with ethanol-induced LORR. While this represents the first time the DO mice have been characterized for level of response to ethanol, others have studied the DO and related populations for additional ethanol-related traits. For example, Hitzemann et al (Hitzemann et al. 2020) recently examined both chronic ethanol consumption and gene expression in the Heterogeneous Stock Collaborative Cross mice (HS-CC, derived from the same eight founders as DO); Bagley et al (Bagley et al. 2021) measured heritability of ethanol consumption and pharmacokinetics in CC strains, and we are aware of ongoing work by Dr. Michael Miles to identify genes associated with ethanol consumption in DO mice. Collectively, these studies demonstrate the ways in the high genetic and phenotypic variation captured by these complementary mouse resources can be used to map genes and identify gene expression networks associated with ethanol-relevant traits. Each QTL accounted for ∼4-5% of the variance explained. As predicted in power simulations, 500 DO mice provide ∼45% power to identify QTLs that explain 5% of the phenotypic variance (Gatti et al 2014). Thus, an even larger sample size than ours (n = 798) would likely increase our ability to detect QTLs of small effect. In addition, future work should include female mice to identify sex-specific QTLs associated with ethanol sensitivity. Despite these limitations, which can be easily addressed in this extendable population, we were able to identify QTLs and obtain favorable mapping resolution. Based on our current results, we predict that a minimum sample size of 1,000 DO mice is appropriate for genetic mapping of most behavioral traits with moderate heritability; even larger sample sizes are expected to produce exponentially larger numbers of significant results (Chitre et al. 2020).

### Gene expression-trait correlations

We also measured gene expression in a subset of the DO mice and performed gene expression-trait correlations to parse amongst positional candidates. We then queried MGI, the NGHRI-GWAS catalog, PubMed, and GeneWeaver to search for associations between these genes and ethanol-relevant traits. Two of the three striatally-expressed genes that we identified were members of the protocadherin gamma subfamily (*Pcdhga7* and *Pcdhgb4*) and showed negative correlations with ethanol-induced ataxia.

### eQTL mapping

Next, we leveraged an existing striatal gene expression dataset from a separate cohort of ethanol naïve DO mice to map eQTLs that co-mapped with the behavioral QTLs identified in our population. This allowed us to further prioritize among the list of candidate genes by examining whether those genes with eQTLs contained SNPs in linkage disequilibrium with the peak marker of the behavioral QTL and by testing the correlation between the expression patterns of each gene to its haplotype effects for the QTL (Table 2). We then queried MGI, the NGHRI-GWAS catalog, and PubMed to search for associations between these genes and ethanol-relevant traits. Our results demonstrate the advantages of integrating high-resolution mapping of behavioral QTLs and eQTLs. Many of the genes we identified in this manner have previously been associated with ethanol phenotypes in rodents or humans, but some were not. Genes in the latter category may provide novel targets for functional validation in follow-up studies.

For the ethanol-induced ataxia QTL on chromosome 16, 19 of the 61 genes in the interval had significant eQTLs. Ten of those genes possessed SNPs with *D′* > .8 (*Cep97, Tbc1d23, Impg2, Nfkbiz, Abi3bp, Cldnd1, Riox2, Cpox, Senp7, Pcnp*) and three (*Cpox, R* = .72; *Arl6, R* = .74; and *Riox2, R* = .72) showed expression patterns that were significantly correlated with the founder haplotype effects. Some of these genes have been associated with alcohol-related phenotypes in humans and mice. For example, a SNP within ABI3BP showed a suggestive (rs12632235, 5 x 10^-6^) association with alcohol dependence in ∼7k Dutch subjects (Mbarek et al. 2015b). Serum levels of ARL6 have been proposed as a novel biomarker for the detection of excessive alcohol use in humans (Liangpunsakul et al. 2015). SNPs within SENP7 showed a significant (rs142338804, 1 x 10^-8^) association with drinks per week in a combined sample of ∼1 million individuals of European ancestry (Karlsson Linnér et al. 2019) and a suggestive association (rs145898511, 8 x 10^-6^) with bitter alcohol consumption (Zhong et al. 2019) in ∼370k subjects of European ancestry. The expression of *Cpox* in whole brain is correlated with ethanol withdrawal severity in BXD RI mice (GS430; (Buck et al. 1997)), and multiple studies have shown differential expression of CPOX in the frontal cortex and hippocampus between alcoholic and nonalcoholic individuals (GS37108; GS246373; GS246393; (Liu et al. 2006; Zhou et al. 2011; Farris et al. 2015)).

For the hypothermia QTL on chromosome 1, 26 of the 72 genes in the interval had significant eQTLs, but only 14 contained SNPs with *D*′ > .8 (*Phlda3, Gm37333, Lmod1, Ipo9, Ptpn7, Ppp1r12b, Nav1, Gm26781, Arl8a, Rnpep, Zbed6, Zfp281, Gm37949, Chil1*); and five in particular (*Nav1, R* = -.71; *Lmod1, R* = -.94; *Klhl12, R* = -.74; *Arl8a, R* = -.85; and *Rnpep, R* = -.73) showed haplotype expression patterns that were significantly correlated with the haplotype behavioral effects. Of these genes, there are a number of interesting candidates. For example, *Phlda3, Ppp1r12b,* and *Ptpn7* have been shown to be differentially expressed in the central amygdala of heterogeneous stock-collaborative cross mice selected for ethanol preference (Colville et al. 2018). *Nav1* is upregulated in response to ethanol and involved in alcohol-induced neuroinflammation in mice (Ureña-Peralta et al. 2018; Xu et al. 2019). ARL8A and KLHL12 have been shown to be differentially expressed in the hippocampus in humans with alcoholism and in humans addicted to cocaine (GS246373, GS246374; (Farris et al. 2015). In addition, the expression of *Arl8a* is significantly altered in the prefrontal cortex of mice four hours after an acute dose of ethanol (GS73824; (Wolen et al. 2012)). Finally, whole brain gene expression of *Rnpep* correlates with blood ethanol concentrations in BXD mice (GS33996; (Philip et al. 2010)), its expression in dopaminergic neurons in the ventral tegmental area is correlated with alcohol consumption (GS246644; (Marballi et al. 2016)), and *Rnpep* is one of 16 preferentially expressed genes in inbred short-sleep mice as compared to their long-sleep counterparts (GS1264; (Xu et al. 2001)).

For the LORR QTL, 37 of the 69 genes in the interval had significant eQTLs that co-mapped with the behavioral QTL on chromosome 2. Among those, 12 genes contained SNPs with *D*′ > .8 (*Fbln7, Anapc1, Mertk, Gm10762, Zc3h8, Fahd2a, Zfp661, Pdyn, Sppl2a, Gm14212, Eid1, Cep152*), and seven genes (*Slc27a2, R* = .94; *Sppl2a, R* = .76; *Anapc1, R* = .73; *Mertk, R* = .75; *Fahd2a, R* = .77; *Cep152, R* = -.71; and *Mrps5, R* = .79) displayed expression patterns were significantly correlated with the founder haplotype effects. Many of these genes have previously been associated with ethanol-relevant traits in mice and/or humans. For example, *Anapc1* is differentially expressed in the brains of mice selectively bred for differences in acute functional tolerance to an incoordinating effect of ethanol as measured by LORR duration (GS1777; (Tabakoff et al. 2003)), its expression in the neocortex is correlated with handling induced convulsions following ethanol administration in BXD mice (GS35918; GS35936; GS35960; GS35964; (Philip et al. 2010, p. 2010)), and ANAPC1 has been shown to be differentially expressed in the basolateral amygdala and in the central nucleus of the amygdala in humans with alcoholism as compared to controls (GS137407; GS137413; (Ponomarev et al. 2012)). The expression of *Mertk* and *Cep152* was shown to be significantly and consistently altered in a meta-analysis comprised of three selected lines and six isogenic strains of mice known to differ in voluntary alcohol consumption (GS3647; (Mulligan et al. 2006)). In addition, *Mertk* showed differential expression in the nucleus accumbens of ethanol drinking in female mice selectively bred to drink to intoxication (HDID) that were treated with vehicle as compared to the water drinking and vehicle treated control group (GS356468; (Pozhidayeva et al. 2020)). Expression levels of *Sppl2a* in the prefrontal cortex of BXD mice were significantly changed following an acute dose of ethanol (GS73824; (Wolen et al. 2012)) and SPPL2a was shown to be differentially expressed in the hippocampus of alcohol and cocaine abusers as compared to controls (GS246375; (Farris et al. 2015)). Female mice lacking *Pdyn* showed significantly lower preference and consumption of ethanol, but interestingly, no differences in LORR (Blednov et al. 2006). In humans, multiple studies have demonstrated that PDYN haplotypes and individual SNPs (rs2281285) are associated with alcohol dependence and propensity to drink in negative emotional states (Flory et al. 2011; Karpyak et al. 2013; Winham et al. 2015), and that decreased methylation levels of PDYN are associated with craving and alcohol consumption (D’Addario et al. 2017).

Given that the majority of human GWAS findings implicate regulatory rather than coding differences (Albert and Kruglyak 2015; Markunas et al. 2017), eQTL mapping can help clarify the link between a genomic region implicated by GWAS and the biological processes that underlie it. However, eQTL mapping is most effective when the tissue of expression is relevant to the trait of interest; and in some cases, the same regulatory region and variant may be linked to one gene in one tissue and another gene in another tissue (Nica and Dermitzakis 2013). Although the striatum is certainly a brain region that is relevant to AUDs broadly, future studies should examine other tissues (such as motor cortex, hypothalamus, and cerebellum) that are known to be involved in ethanol-induced ataxia, hypothermia, and LORR, to ensure detection of trait relevant eQTLs. It is also possible that eQTLs are present at an earlier developmental time-point, at a different time-period post-ethanol exposure, or in a small minority of cells, in which case our study design would fail to detect them.

### GeneWeaver

Lastly, we used GeneWeaver to identify highly connected genes across datasets and examined their relevance to alcohol-related traits using bioinformatics databases. For ethanol-induced ataxia, we identified four genes (*Tomm70a, Pcnp, Abi3bp* and *Trmt10c*) shared across multiple gene-sets. *Abi3bp*, *Tomm70a,* and *Trmt10c* have been linked to ethanol-relevant phenotypes. As described earlier, a SNP within ABI3BP showed a suggestive (rs12632235, 5 x 10^-6^) association with alcohol dependence in ∼7k Dutch subjects (Mbarek et al. 2015). The expression of *Tomm70* in the nucleus accumbens, prefrontal cortex, and ventral tegmental area has been shown to differ in C57BL/6J and DBA/2J mice four hours after ethanol exposure (GS1547; (Kerns et al. 2005)), and in the striatum and cortex of C57BL/6J mice drinking to intoxication (GS127341, GS127342;(Mulligan et al. 2011)). Lastly, *Trmt10* is differentially expressed in the nucleus accumbens following continuous alcohol drinking in alcohol preferring rats (GS37147; (Bell et al. 2009). Although *Pcnp* has not been previously associated with alcohol-related phenotypes, it is known to be involved in protein ubiquitination. Given that alcohol exposure significantly alters many post-translational processes including ubiquitination, *Pcnp* may represent an ethanol-responsive gene that is involved in a network of neural adaptations that lead to dependence (Wolen et al. 2012).

For ethanol-induced hypothermia, we identified eight genes shared between various gene-sets (*Klhl12, Tnnt2, Adora1, 9230116N13Rik, Mybph, Lamc2, Dpp10,* and *Creg2*), with *Klhl12* representing the most highly connected gene we identified via GeneWeaver. As described earlier, *KLHL12* is differentially expressed in the hippocampus of humans with alcoholism and in humans with cocaine addiction (Farris et al. 2015). Another gene, *Adora1,* has been previously linked to ethanol phenotypes in both mice and humans. Mice lacking *Adora1* display abnormal body temperature homeostasis (J:70664; (Johansson et al. 2001), numerous groups have shown that *Adora1* is differentially expressed in the brains of mice and rats that differ in ethanol consumption or preference ((Colville et al. 2018); GS3647; (Mulligan et al. 2008); GS31782; (Phillips et al. 1994); GS75752; (Worst et al. 2005)), and adenosine A1 receptors are required for astrocyte calcium activation to increase ethanol sedation (Erickson et al. 2020). Lastly, a variant set associated with ADORA1 showed a suggestive association (5.29 x 10^-5^) with alcohol dependence in 806 European Americans (Clark et al. 2017). Another highly connected gene was *Tnnt2*. In a sample of 8754 individuals from the Australian Twin Registry, SNPs within TNNT2 were nominally associated (7.4 x 10^-5^) with weekly alcohol consumption over a 1-year period (Heath et al. 2011). In addition, *Tnnt2* has been shown to be differentially expressed in response to alcohol in the mouse anterior cingulate cortex (GS246728; Repunte-Canonigo et al., 2010). Expression of *9230116N13Rik* and *Mybph* in the neocortex and striatum respectively, has been correlated with handling induced convulsions following ethanol in the BXD RI panel (GS35908, GS35902; (Philip et al. 2010). In humans, a SNP within LAMC2 was associated (rs1047980; 9 x 10^-6^) with maximum habitual alcohol consumption ∼140,000 U.S. European and African American veterans (Gelernter et al. 2019a). In addition, *Lamc2* shows differential expression in the nucleus accumbens of alcohol preferring vs non-preferring rats (GS246645, (Stankiewicz et al. 2015)) and its expression in the hippocampus correlates with handling induced convulsions in BXD mice (GS35963, (Philip et al. 2010)). DPP10 is differentially expressed in the hippocampus of human cocaine addicts (GS246374; (Farris et al. 2015)), and *Dpp10* shows changes in gene expression in dopaminergic neurons in the ventral tegmental area following binge-like alcohol consumption in alcohol preferring mice (GS246644; (Marballi et al. 2016)). Lastly, expression levels of *Creg2* in the prefrontal cortex were significantly altered four hours after an acute dose of ethanol in BXD RI mice (GS73824; (Wolen et al. 2012)), and it shows differential expression in the nucleus accumbens in mice that were selectively bred to drink to intoxication (GS356469, (Pozhidayeva et al. 2020)).

Finally, for ethanol-induced LORR, we identified three genes (*Prkcq, Bub1,* and *Itpripl1*) that were shared across multiple gene-sets. Neither *Prkcq* nor *Bub1* have been linked to ethanol-relevant traits, but *Itpripl1* is differentially expressed in the periaqueductal gray following binge-like alcohol consumption in rats that were selectively bred to prefer alcohol (GS246761, (McClintick et al. 2016)).

## Conclusions

A significant challenge in identifying genes associated with risk for AUD is that most large-scale GWAS have focused primarily on AUD diagnosis or self-reported problematic use (Gelernter et al. 2014; Quillen et al. 2014; Mbarek et al. 2015b; Kranzler et al. 2019; Sanchez-Roige et al. 2019a, p., b, p.; Zhou et al. 2020), but see (Meyers et al. 2016, 2017, 2020; Gelernter et al. 2019b; Lai et al. 2020b; Sanchez-Roige and Palmer 2020). AUD is a complex disease that represents the end point of multiple stages encompassing initial sensitivity to the effects of alcohol, the transition to problematic alcohol use and loss of control, the development of tolerance, and relapse (Sanchez-Roige and Palmer 2020). Each of these stages is likely to be influenced by unique networks of genes. The reliance on broad, binary clinical diagnoses limits the dissection of the genetic architecture of AUD and highlights the need for complementary resources and approaches to overcome the challenges faced by human studies. While there is a large body of literature in humans supporting the relationship between genetically mediated variation in sensitivity to the intoxicating effects of alcohol and later development of AUD (Newlin and Thomson 1990b; Schuckit 1994; Schuckit and Smith 2000, 2011; King et al. 2011; Ramchandani et al. 2011; Schuckit et al. 2014); there are no laboratory-based alcohol challenge GWAS in humans, nor would it be practical to perform one that is sufficiently powered to allow for gene identification. A second obstacle facing human AUD GWAS is the difficulty in identifying the causal genes. One reason for this is due to the fact that most associations implicate non-coding variants that presumably influence risk for AUD by altering gene regulation (Albert and Kruglyak 2015; Markunas et al. 2017). While it has been reliably demonstrated that the use of transcriptomic data in conjunction with systems genetic approaches such as eQTL mapping and bioinformatics network-based analyses can successfully lead to gene identification (Nadeau and Dudley 2011; Civelek and Lusis 2014), few transcriptomic datasets exist for relevant brain tissues in sufficiently large cohorts of individuals with AUDs, thereby limiting the use of systems genetics approaches in AUD GWAS. The goal of the present study was to illustrate the ways in which applying systems genetics approaches to the highly recombinant and genetically diverse DO mice can help address these limitations faced by human AUD GWAS. To accomplish this, we first characterized the DO population for variation in three ethanol-sensitivity endophenotypes and established that they are a valid resource for exploring sensitivity to the ataxic, hypothermic, and intoxicating effects of ethanol. Next, we demonstrated the utility of DO mice for high-resolution mapping of QTLs underlying response to ethanol. We mapped eQTLs, performed expression-trait correlations, and used a series of bioinformatics approaches to integrate results across endophenotypes, expression data, and species in order to parse among the positional candidate genes within our QTL intervals. The data presented here can be used to 1) prioritize human GWAS hits for follow-up investigation, 2) provide additional insight into biological and behavioral mechanisms related to AUD, and 3) enhance our understanding of how these variants influence risk for the development of AUD, something that would be very difficult to obtain using human GWAS alone. Collectively, our results highlight the power of systems genetics in the DO population and demonstrate the ways in which translational mouse research can inform our understanding of the genetic underpinnings of AUD in humans.

## Acknowledgements

This work was supported by a NARSAD BBRF Young Investigator Award (CCP), NIH P20-GM-103449 (CCP), and by the NIAAA Intramural Research Program (AH). EJC is supported by R01DA037937 and P50DA039841. The GeneWeaver software system is supported by AA018776. AAP was supported by P50DA037844 and R01AA026281.

## Reference List

Albert FW, Kruglyak L (2015) The role of regulatory variation in complex traits and disease. Nature Reviews Genetics 16:197–212. https://doi.org/10.1038/nrg3891

Bagley JR, Chesler EJ, Philip VM, et al (2021) Heritability of ethanol consumption and pharmacokinetics in a genetically diverse panel of collaborative cross mouse strains and their inbred founders. Alcohol Clin Exp Res 45:697–708. https://doi.org/10.1111/acer.14582

Bell RL, Kimpel MW, McClintick JN, et al (2009) Gene expression changes in the nucleus accumbens of alcohol-preferring rats following chronic ethanol consumption. Pharmacol Biochem Behav 94:131–147. https://doi.org/10.1016/j.pbb.2009.07.019

Blednov YA, Walker D, Martinez M, Harris RA (2006) Reduced alcohol consumption in mice lacking preprodynorphin. Alcohol 40:73–86. https://doi.org/10.1016/j.alcohol.2006.12.002

Broman KW, Gatti DM, Simecek P, et al (2019) R/qtl2: Software for Mapping Quantitative Trait Loci with High-Dimensional Data and Multiparent Populations. Genetics 211:495–502. https://doi.org/10.1534/genetics.118.301595

Buck KJ, Metten P, Belknap JK, Crabbe JC (1997) Quantitative trait loci involved in genetic predisposition to acute alcohol withdrawal in mice. J Neurosci 17:3946–3955

Cheng R, Parker CC, Abney M, Palmer AA (2013) Practical considerations regarding the use of genotype and pedigree data to model relatedness in the context of genome-wide association studies. G3 (Bethesda) 3:1861–1867. https://doi.org/10.1534/g3.113.007948

Chesler EJ, Plitt A, Fisher D, et al (2012) Quantitative trait loci for sensitivity to ethanol intoxication in a C57BL/6J×129S1/SvImJ inbred mouse cross. Mammalian Genome: Official Journal of the International Mammalian Genome Society 23:305–321. https://doi.org/10.1007/s00335-012-9394-2

Chitre AS, Polesskaya O, Holl K, et al (2020) Genome-Wide Association Study in 3,173 Outbred Rats Identifies Multiple Loci for Body Weight, Adiposity, and Fasting Glucose. Obesity 28:1964–1973. https://doi.org/10.1002/oby.22927

Christensen SC, Johnson TE, Markel PD, et al (1996) Quantitative trait locus analyses of sleep-times induced by sedative-hypnotics in LSXSS recombinant inbred strains of mice. Alcohol Clin Exp Res 20:543–550. https://doi.org/10.1111/j.1530-0277.1996.tb01090.x

Churchill GA, Gatti DM, Munger SC, Svenson KL (2012) The Diversity Outbred mouse population. Mammalian Genome: Official Journal of the International Mammalian Genome Society 23:713– 718. https://doi.org/10.1007/s00335-012-9414-2

Civelek M, Lusis AJ (2014) Systems genetics approaches to understand complex traits. Nat Rev Genet 15:34–48. https://doi.org/10.1038/nrg3575

Clark SL, McClay JL, Adkins DE, et al (2017) Deep Sequencing of 71 Candidate Genes to Characterize Variation Associated with Alcohol Dependence. Alcoholism: Clinical and Experimental Research 41:711–718. https://doi.org/10.1111/acer.13352

Colville AM, Iancu OD, Lockwood DR, et al (2018) Regional Differences and Similarities in the Brain Transcriptome for Mice Selected for Ethanol Preference From HS-CC Founders. Front Genet 9:300. https://doi.org/10.3389/fgene.2018.00300

Crabbe JC, Cameron AJ, Munn E, et al (2008) Overview of mouse assays of ethanol intoxication. Current Protocols in Neuroscience Chapter 9:Unit 9.26. https://doi.org/10.1002/0471142301.ns0926s42

Crabbe JC, Feller DJ, Dorow JS (1989) Sensitivity and tolerance to ethanol-induced hypothermia in genetically selected mice. J Pharmacol Exp Ther 249:456–461

Crabbe JC, Gallaher ES, Phillips TJ, Belknap JK (1994) Genetic determinants of sensitivity to ethanol in inbred mice. Behav Neurosci 108:186–195. https://doi.org/10.1037//0735-7044.108.1.186

D’Addario C, Shchetynsky K, Pucci M, et al (2017) Genetic variation and epigenetic modification of the prodynorphin gene in peripheral blood cells in alcoholism. Prog Neuropsychopharmacol Biol Psychiatry 76:195–203. https://doi.org/10.1016/j.pnpbp.2017.03.012

DuBose CS, Chesler EJ, Goldowitz D, Hamre KM (2013) Use of the expanded panel of BXD mice narrow QTL regions in ethanol-induced locomotor activation and motor incoordination. Alcoholism, Clinical and Experimental Research 37:170–183. https://doi.org/10.1111/j.1530-0277.2012.01865.x

Erickson EK, DaCosta AJ, Mason SC, et al (2020) Cortical astrocytes regulate ethanol consumption and intoxication in mice. Neuropsychopharmacology. https://doi.org/10.1038/s41386-020-0721-0

Farris SP, Harris RA, Ponomarev I (2015) Epigenetic modulation of brain gene networks for cocaine and alcohol abuse. Front Neurosci 9:176. https://doi.org/10.3389/fnins.2015.00176

Flory JD, Pytte CL, Hurd Y, et al (2011) Alcohol dependence, disinhibited behavior and variation in the prodynorphin gene. Biol Psychol 88:51–56. https://doi.org/10.1016/j.biopsycho.2011.06.007

Gatti DM, Svenson KL, Shabalin A, et al (2014) Quantitative trait locus mapping methods for diversity outbred mice. G3 (Bethesda, Md) 4:1623–1633. https://doi.org/10.1534/g3.114.013748

Gehle VM, Erwin VG (2000) The genetics of acute functional tolerance and initial sensitivity to ethanol for an ataxia test in the LSxSS RI strains. Alcoholism, Clinical and Experimental Research 24:579–587

Gelernter J, Kranzler HR, Sherva R, et al (2014) Genome-wide association study of alcohol dependence:significant findings in African-and European-Americans including novel risk loci. Molecular Psychiatry 19:41–49. https://doi.org/10.1038/mp.2013.145

Gelernter J, Sun N, Polimanti R, et al (2019a) Genome-wide Association Study of Maximum Habitual Alcohol Intake in >140,000 U.S. European and African American Veterans Yields Novel Risk Loci. Biol Psychiatry 86:365–376. https://doi.org/10.1016/j.biopsych.2019.03.984

Gelernter J, Sun N, Polimanti R, et al (2019b) Genome-wide Association Study of Maximum Habitual Alcohol Intake in >140,000 U.S. European and African American Veterans Yields Novel Risk Loci. Biol Psychiatry 86:365–376. https://doi.org/10.1016/j.biopsych.2019.03.984

Grant BF (1998) The impact of a family history of alcoholism on the relationship between age at onset of alcohol use and DSM-IV alcohol dependence: results from the National Longitudinal Alcohol Epidemiologic Survey. Alcohol Health and Research World 22:144–147

Hart AB, Kranzler HR (2015) Alcohol Dependence Genetics: Lessons Learned From Genome-Wide Association Studies (GWAS) and Post-GWAS Analyses. Alcoholism, Clinical and Experimental Research 39:1312–1327. https://doi.org/10.1111/acer.12792

Heath AC, Whitfield JB, Martin NG, et al (2011) A quantitative-trait genome-wide association study of alcoholism risk in the community: findings and implications. Biol Psychiatry 70:513–518. https://doi.org/10.1016/j.biopsych.2011.02.028

Hindorff LA, Sethupathy P, Junkins HA, et al (2009) Potential etiologic and functional implications of genome-wide association loci for human diseases and traits. Proceedings of the National Academy of Sciences of the United States of America 106:9362–9367. https://doi.org/10.1073/pnas.0903103106

Hitzemann R, Phillips TJ, Lockwood DR, et al (2020) Phenotypic and gene expression features associated with variation in chronic ethanol consumption in heterogeneous stock collaborative cross mice. Genomics. https://doi.org/10.1016/j.ygeno.2020.08.004

Holdstock L, King AC, de Wit H (2000) Subjective and objective responses to ethanol in moderate/heavy and light social drinkers. Alcoholism, Clinical and Experimental Research 24:789–794

Johansson B, Halldner L, Dunwiddie TV, et al (2001) Hyperalgesia, anxiety, and decreased hypoxic neuroprotection in mice lacking the adenosine A1 receptor. PNAS 98:9407–9412. https://doi.org/10.1073/pnas.161292398

Karlsson Linnér R, Biroli P, Kong E, et al (2019) Genome-wide association analyses of risk tolerance and risky behaviors in over 1 million individuals identify hundreds of loci and shared genetic influences. Nat Genet 51:245–257. https://doi.org/10.1038/s41588-018-0309-3

Karpyak VM, Winham SJ, Preuss UW, et al (2013) Association of the PDYN gene with alcohol dependence and the propensity to drink in negative emotional states. Int J Neuropsychopharmacol 16:975–985. https://doi.org/10.1017/S1461145712001137

Keane TM, Goodstadt L, Danecek P, et al (2011) Mouse genomic variation and its effect on phenotypes and gene regulation. Nature 477:289–294. https://doi.org/10.1038/nature10413

Kendler KS, Aggen SH, Prescott CA, et al (2012) Evidence for multiple genetic factors underlying the DSM-IV criteria for alcohol dependence. Mol Psychiatry 17:1306–1315. https://doi.org/10.1038/mp.2011.153

Kerns RT, Ravindranathan A, Hassan S, et al (2005) Ethanol-responsive brain region expression networks: implications for behavioral responses to acute ethanol in DBA/2J versus C57BL/6J mice. J Neurosci 25:2255–2266. https://doi.org/10.1523/JNEUROSCI.4372-04.2005

King AC, de Wit H, McNamara PJ, Cao D (2011) Rewarding, stimulant, and sedative alcohol responses and relationship to future binge drinking. Archives of General Psychiatry 68:389–399. https://doi.org/10.1001/archgenpsychiatry.2011.26

Kranzler HR, Zhou H, Kember RL, et al (2019) Genome-wide association study of alcohol consumption and use disorder in 274,424 individuals from multiple populations. Nat Commun 10:1499. https://doi.org/10.1038/s41467-019-09480-8

Krystal JH, Petrakis IL, Mason G, et al (2003) N-methyl-D-aspartate glutamate receptors and alcoholism: reward, dependence, treatment, and vulnerability. Pharmacol Ther 99:79–94

Lai D, Kapoor M, Wetherill L, et al (2020a) Genome-wide admixture mapping of DSM-IV alcohol dependence, criterion count, and the self-rating of the effects of ethanol in African American populations. Am J Med Genet B Neuropsychiatr Genet. https://doi.org/10.1002/ajmg.b.32805

Lai D, Wetherill L, Kapoor M, et al (2020b) Genome-wide association studies of the self-rating of effects of ethanol (SRE). Addict Biol 25:e12800. https://doi.org/10.1111/adb.12800

Liangpunsakul S, Lai X, Ross RA, et al (2015) Novel serum biomarkers for detection of excessive alcohol use. Alcohol Clin Exp Res 39:556–565. https://doi.org/10.1111/acer.12654

Liu J, Lewohl JM, Harris RA, et al (2006) Patterns of gene expression in the frontal cortex discriminate alcoholic from nonalcoholic individuals. Neuropsychopharmacology 31:1574–1582. https://doi.org/10.1038/sj.npp.1300947

Liu M, Jiang Y, Wedow R, et al (2019) Association studies of up to 1.2 million individuals yield new insights into the genetic etiology of tobacco and alcohol use. Nat Genet 51:237–244. https://doi.org/10.1038/s41588-018-0307-5

Marballi K, Genabai NK, Blednov YA, et al (2016) Alcohol consumption induces global gene expression changes in VTA dopaminergic neurons. Genes Brain Behav 15:318–326. https://doi.org/10.1111/gbb.12266

Markel PD, Bennett B, Beeson M, et al (1997) Confirmation of quantitative trait loci for ethanol sensitivity in long-sleep and short-sleep mice. Genome Res 7:92–99. https://doi.org/10.1101/gr.7.2.92

Markunas CA, Johnson EO, Hancock DB (2017) Comprehensive evaluation of disease- and trait-specific enrichment for eight functional elements among GWAS-identified variants. Hum Genet 136:911– 919. https://doi.org/10.1007/s00439-017-1815-6

Mbarek H, Milaneschi Y, Fedko IO, et al (2015a) The genetics of alcohol dependence: Twin and SNP-based heritability, and genome-wide association study based on AUDIT scores. Am J Med Genet B Neuropsychiatr Genet 168:739–748. https://doi.org/10.1002/ajmg.b.32379

Mbarek H, Milaneschi Y, Fedko IO, et al (2015b) The genetics of alcohol dependence: Twin and SNP-based heritability, and genome-wide association study based on AUDIT scores. American Journal of Medical Genetics Part B: Neuropsychiatric Genetics 168:739–748. https://doi.org/10.1002/ajmg.b.32379

McClintick JN, McBride WJ, Bell RL, et al (2016) Gene Expression Changes in Glutamate and GABA-A Receptors, Neuropeptides, Ion Channels, and Cholesterol Synthesis in the Periaqueductal Gray Following Binge-Like Alcohol Drinking by Adolescent Alcohol-Preferring (P) Rats. Alcohol Clin Exp Res 40:955–968. https://doi.org/10.1111/acer.13056

Meyers JL, Zhang J, Chorlian DB, et al (2020) A genome-wide association study of interhemispheric theta EEG coherence: implications for neural connectivity and alcohol use behavior. Mol Psychiatry. https://doi.org/10.1038/s41380-020-0777-6

Meyers JL, Zhang J, Manz N, et al (2016) A Genome Wide Association Study of Fast Beta EEG in Families of European Ancestry. Int J Psychophysiol. https://doi.org/10.1016/j.ijpsycho.2016.12.008

Meyers JL, Zhang J, Wang JC, et al (2017) An endophenotype approach to the genetics of alcohol dependence: a genome wide association study of fast beta EEG in families of African ancestry. Mol Psychiatry. https://doi.org/10.1038/mp.2016.239

Morgan AP, Fu C-P, Kao C-Y, et al (2015) The Mouse Universal Genotyping Array: From Substrains to Subspecies. G3 (Bethesda, Md) 6:263–279. https://doi.org/10.1534/g3.115.022087

Mulligan MK, Ponomarev I, Boehm SL, et al (2008) Alcohol trait and transcriptional genomic analysis of C57BL/6 substrains. Genes Brain Behav 7:677–689. https://doi.org/10.1111/j.1601-183X.2008.00405.x

Mulligan MK, Ponomarev I, Hitzemann RJ, et al (2006) Toward understanding the genetics of alcohol drinking through transcriptome meta-analysis. Proceedings of the National Academy of Sciences of the United States of America 103:6368–6373. https://doi.org/10.1073/pnas.0510188103

Mulligan MK, Rhodes JS, Crabbe JC, et al (2011) Molecular profiles of drinking alcohol to intoxication in C57BL/6J mice. Alcohol Clin Exp Res 35:659–670. https://doi.org/10.1111/j.1530-0277.2010.01384.x

Nadeau JH, Dudley AM (2011) Genetics. Systems genetics. Science 331:1015–1016. https://doi.org/10.1126/science.1203869

Newlin DB, Thomson JB (1990a) Alcohol challenge with sons of alcoholics: a critical review and analysis. Psychological Bulletin 108:383–402

Newlin DB, Thomson JB (1990b) Alcohol challenge with sons of alcoholics: a critical review and analysis. Psychol Bull 108:383–402

Nica AC, Dermitzakis ET (2013) Expression quantitative trait loci: present and future. Philos Trans R Soc Lond B Biol Sci 368:. https://doi.org/10.1098/rstb.2012.0362

Parker CC, Lusk R, Saba LM (2020) Alcohol Sensitivity as an Endophenotype of Alcohol Use Disorder: Exploring Its Translational Utility between Rodents and Humans. Brain Sci 10:. https://doi.org/10.3390/brainsci10100725

Parker CC, Palmer AA (2011) Dark matter: are mice the solution to missing heritability? Frontiers in Genetics 2:32. https://doi.org/10.3389/fgene.2011.00032

Philip VM, Duvvuru S, Gomero B, et al (2010) High-throughput behavioral phenotyping in the expanded panel of BXD recombinant inbred strains. Genes, Brain, and Behavior 9:129–159. https://doi.org/10.1111/j.1601-183X.2009.00540.x

Phillips TJ, Crabbe JC, Metten P, Belknap JK (1994) Localization of genes affecting alcohol drinking in mice. Alcohol Clin Exp Res 18:931–941. https://doi.org/10.1111/j.1530-0277.1994.tb00062.x

Ponomarev I, Wang S, Zhang L, et al (2012) Gene coexpression networks in human brain identify epigenetic modifications in alcohol dependence. J Neurosci 32:1884–1897. https://doi.org/10.1523/JNEUROSCI.3136-11.2012

Pozhidayeva DY, Farris SP, Goeke CM, et al (2020) Chronic Chemogenetic Stimulation of the Nucleus Accumbens Produces Lasting Reductions in Binge Drinking and Ameliorates Alcohol-Related Morphological and Transcriptional Changes. Brain Sci 10:. https://doi.org/10.3390/brainsci10020109

Quillen EE, Chen X-D, Almasy L, et al (2014) ALDH2 is associated to alcohol dependence and is the major genetic determinant of “daily maximum drinks” in a GWAS study of an isolated rural Chinese sample. Am J Med Genet B Neuropsychiatr Genet 165B:103–110. https://doi.org/10.1002/ajmg.b.32213

Quinn PD, Fromme K (2011) Subjective response to alcohol challenge: a quantitative review. Alcohol Clin Exp Res 35:1759–1770. https://doi.org/10.1111/j.1530-0277.2011.01521.x

Ramchandani VA, Umhau J, Pavon FJ, et al (2011) A genetic determinant of the striatal dopamine response to alcohol in men. Mol Psychiatry 16:809–817. https://doi.org/10.1038/mp.2010.56

Repunte-Canonigo V, van der Stap LD, Chen J, et al (2010) Genome-wide gene expression analysis identifies K-ras as a regulator of alcohol intake. Brain Res 1339:1–10. https://doi.org/10.1016/j.brainres.2010.03.063

Sanchez-Roige S, Fontanillas P, Elson SL, et al (2019a) Genome-wide association study of alcohol use disorder identification test (AUDIT) scores in 20 328 research participants of European ancestry. Addict Biol 24:121–131. https://doi.org/10.1111/adb.12574

Sanchez-Roige S, Palmer AA (2020) Emerging phenotyping strategies will advance our understanding of psychiatric genetics. Nature Neuroscience 23:475–480. https://doi.org/10.1038/s41593-020-0609-7

Sanchez-Roige S, Palmer AA, Fontanillas P, et al (2019b) Genome-Wide Association Study Meta-Analysis of the Alcohol Use Disorders Identification Test (AUDIT) in Two Population-Based Cohorts. Am J Psychiatry 176:107–118. https://doi.org/10.1176/appi.ajp.2018.18040369

Saul MC, Philip VM, Reinholdt LG, et al (2019) High-Diversity Mouse Populations for Complex Traits. Trends Genet 35:501–514. https://doi.org/10.1016/j.tig.2019.04.003

Savage JE, Neale Z, Cho SB, et al (2015) Level of response to alcohol as a factor for targeted prevention in college students. Alcohol Clin Exp Res 39:2215–2223. https://doi.org/10.1111/acer.12874

Schuckit MA (1994) Low level of response to alcohol as a predictor of future alcoholism. The American Journal of Psychiatry 151:184–189

Schuckit MA, Smith TL (2000) The relationships of a family history of alcohol dependence, a low level of response to alcohol and six domains of life functioning to the development of alcohol use disorders. Journal of Studies on Alcohol 61:827–835

Schuckit MA, Smith TL (2001) A comparison of correlates of DSM-IV alcohol abuse or dependence among more than 400 sons of alcoholics and controls. Alcoholism, Clinical and Experimental Research 25:1–8

Schuckit MA, Smith TL (2011) Onset and course of alcoholism over 25 years in middle class men. Drug and Alcohol Dependence 113:21–28. https://doi.org/10.1016/j.drugalcdep.2010.06.017

Schuckit MA, Smith TL, Clausen P, et al (2016) The Low Level of Response to Alcohol-Based Heavy Drinking Prevention Program: One-Year Follow-Up. J Stud Alcohol Drugs 77:25–37. https://doi.org/10.15288/jsad.2016.77.25

Schuckit MA, Smith TL, Kalmijn JA (2014) The patterns of drug and alcohol use and associated problems over 30 years in 397 men. Alcohol Clin Exp Res 38:227–234. https://doi.org/10.1111/acer.12220

Stankiewicz AM, Goscik J, Dyr W, et al (2015) Novel candidate genes for alcoholism--transcriptomic analysis of prefrontal medial cortex, hippocampus and nucleus accumbens of Warsaw alcohol-preferring and non-preferring rats. Pharmacol Biochem Behav 139:27–38. https://doi.org/10.1016/j.pbb.2015.10.003

Stranger BE, Stahl EA, Raj T (2011) Progress and promise of genome-wide association studies for human complex trait genetics. Genetics 187:367–383. https://doi.org/10.1534/genetics.110.120907

Svenson KL, Gatti DM, Valdar W, et al (2012) High-resolution genetic mapping using the Mouse Diversity outbred population. Genetics 190:437–447. https://doi.org/10.1534/genetics.111.132597

Tabakoff B, Bhave SV, Hoffman PL (2003) Selective breeding, quantitative trait locus analysis, and gene arrays identify candidate genes for complex drug-related behaviors. J Neurosci 23:4491–4498

Ureña-Peralta JR, Alfonso-Loeches S, Cuesta-Diaz CM, et al (2018) Deep sequencing and miRNA profiles in alcohol-induced neuroinflammation and the TLR4 response in mice cerebral cortex. Sci Rep 8:15913. https://doi.org/10.1038/s41598-018-34277-y

Verhulst B, Neale MC, Kendler KS (2015) The heritability of alcohol use disorders: a meta-analysis of twin and adoption studies. Psychol Med 45:1061–1072. https://doi.org/10.1017/S0033291714002165

Ward LD, Kellis M (2012) Interpreting noncoding genetic variation in complex traits and human disease. Nature Biotechnology 30:1095–1106. https://doi.org/10.1038/nbt.2422

Winham SJ, Preuss UW, Geske JR, et al (2015) Associations of prodynorphin sequence variation with alcohol dependence and related traits are phenotype-specific and sex-dependent. Sci Rep 5:15670. https://doi.org/10.1038/srep15670

Wolen AR, Phillips CA, Langston MA, et al (2012) Genetic dissection of acute ethanol responsive gene networks in prefrontal cortex: functional and mechanistic implications. PloS One 7:e33575. https://doi.org/10.1371/journal.pone.0033575

Worst TJ, Tan JC, Robertson DJ, et al (2005) Transcriptome analysis of frontal cortex in alcohol-preferring and nonpreferring rats. J Neurosci Res 80:529–538. https://doi.org/10.1002/jnr.20496

Xu W, Liyanage VRB, MacAulay A, et al (2019) Genome-Wide Transcriptome Landscape of Embryonic Brain-Derived Neural Stem Cells Exposed to Alcohol with Strain-Specific Cross-Examination in BL6 and CD1 Mice. Scientific Reports 9:206. https://doi.org/10.1038/s41598-018-36059-y

Xu Y, Ehringer M, Yang F, Sikela JM (2001) Comparison of global brain gene expression profiles between inbred long-sleep and inbred short-sleep mice by high-density gene array hybridization. Alcohol Clin Exp Res 25:810–818

Zhong VW, Kuang A, Danning RD, et al (2019) A genome-wide association study of bitter and sweet beverage consumption. Hum Mol Genet 28:2449–2457. https://doi.org/10.1093/hmg/ddz061

Zhou H, Sealock JM, Sanchez-Roige S, et al (2020) Genome-wide meta-analysis of problematic alcohol use in 435,563 individuals yields insights into biology and relationships with other traits. Nature Neuroscience 1–10. https://doi.org/10.1038/s41593-020-0643-5

Zhou Z, Yuan Q, Mash DC, Goldman D (2011) Substance-specific and shared transcription and epigenetic changes in the human hippocampus chronically exposed to cocaine and alcohol. Proc Natl Acad Sci USA 108:6626–6631. https://doi.org/10.1073/pnas.1018514108

