## Supplementary Figures for "Genome-wide association mapping of ethanol sensitivity in the Diversity Outbred mouse population"

### Slide 1
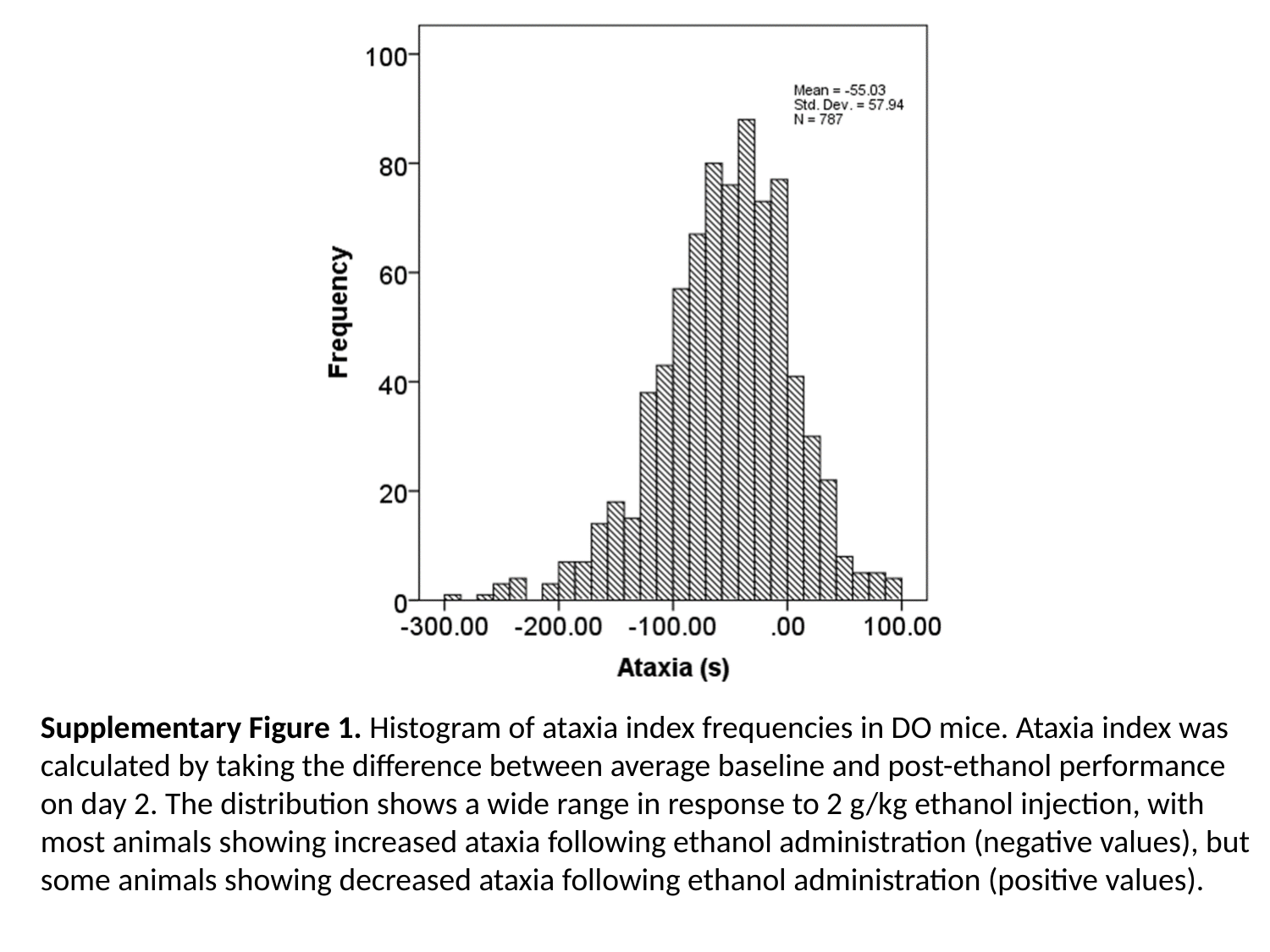

Supplementary Figure 1. Histogram of ataxia index frequencies in DO mice. Ataxia index was calculated by taking the difference between average baseline and post-ethanol performance on day 2. The distribution shows a wide range in response to 2 g/kg ethanol injection, with most animals showing increased ataxia following ethanol administration (negative values), but some animals showing decreased ataxia following ethanol administration (positive values).

### Slide 2
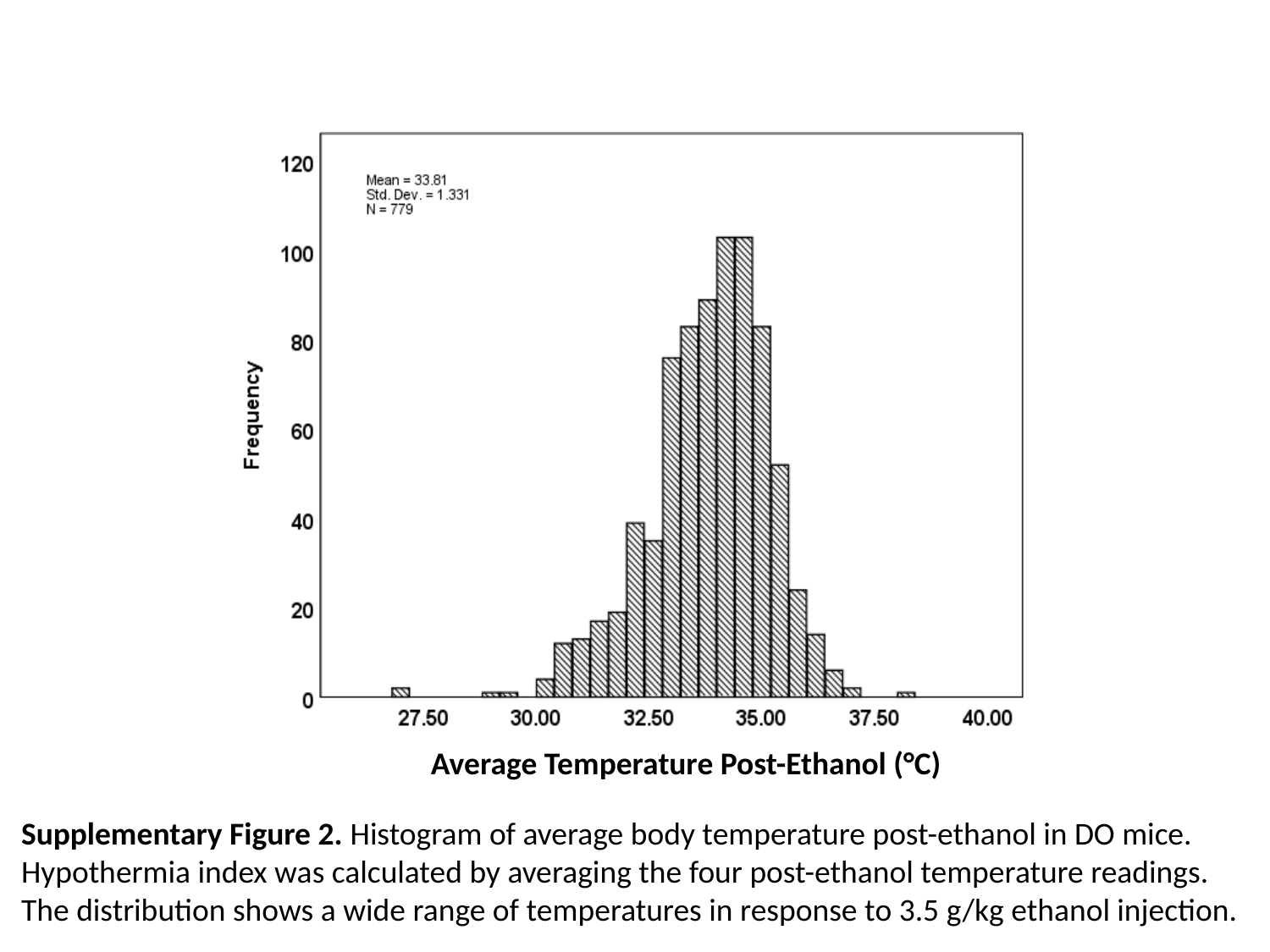

Average Temperature Post-Ethanol (°C)
Supplementary Figure 2. Histogram of average body temperature post-ethanol in DO mice. Hypothermia index was calculated by averaging the four post-ethanol temperature readings. The distribution shows a wide range of temperatures in response to 3.5 g/kg ethanol injection.

### Slide 3
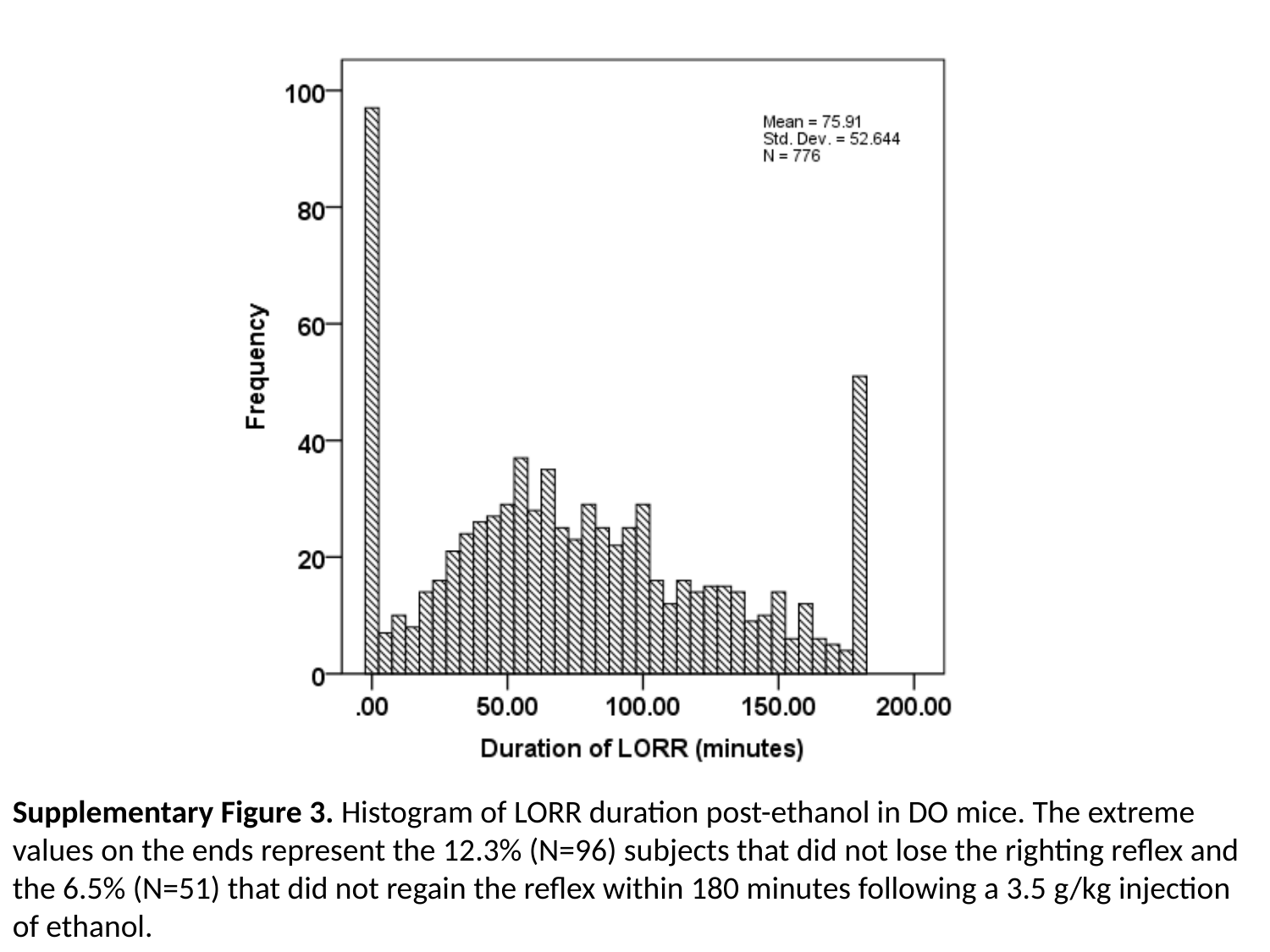

Supplementary Figure 3. Histogram of LORR duration post-ethanol in DO mice. The extreme values on the ends represent the 12.3% (N=96) subjects that did not lose the righting reflex and the 6.5% (N=51) that did not regain the reflex within 180 minutes following a 3.5 g/kg injection of ethanol.

### Slide 4
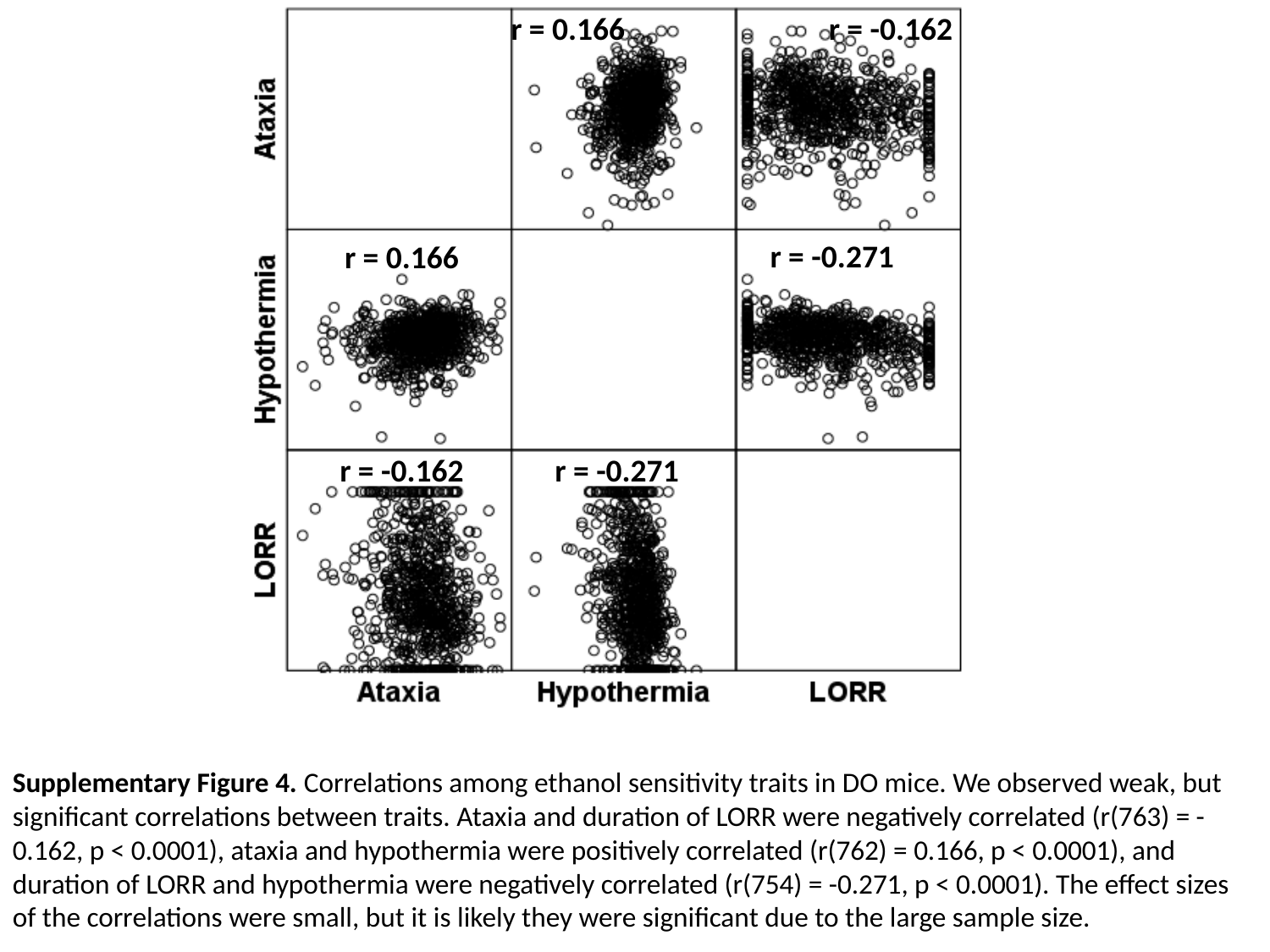

r = 0.166
r = -0.162
r = -0.271
r = 0.166
r = -0.162
r = -0.271
Supplementary Figure 4. Correlations among ethanol sensitivity traits in DO mice. We observed weak, but significant correlations between traits. Ataxia and duration of LORR were negatively correlated (r(763) = -0.162, p < 0.0001), ataxia and hypothermia were positively correlated (r(762) = 0.166, p < 0.0001), and duration of LORR and hypothermia were negatively correlated (r(754) = -0.271, p < 0.0001). The effect sizes of the correlations were small, but it is likely they were significant due to the large sample size.

### Slide 5
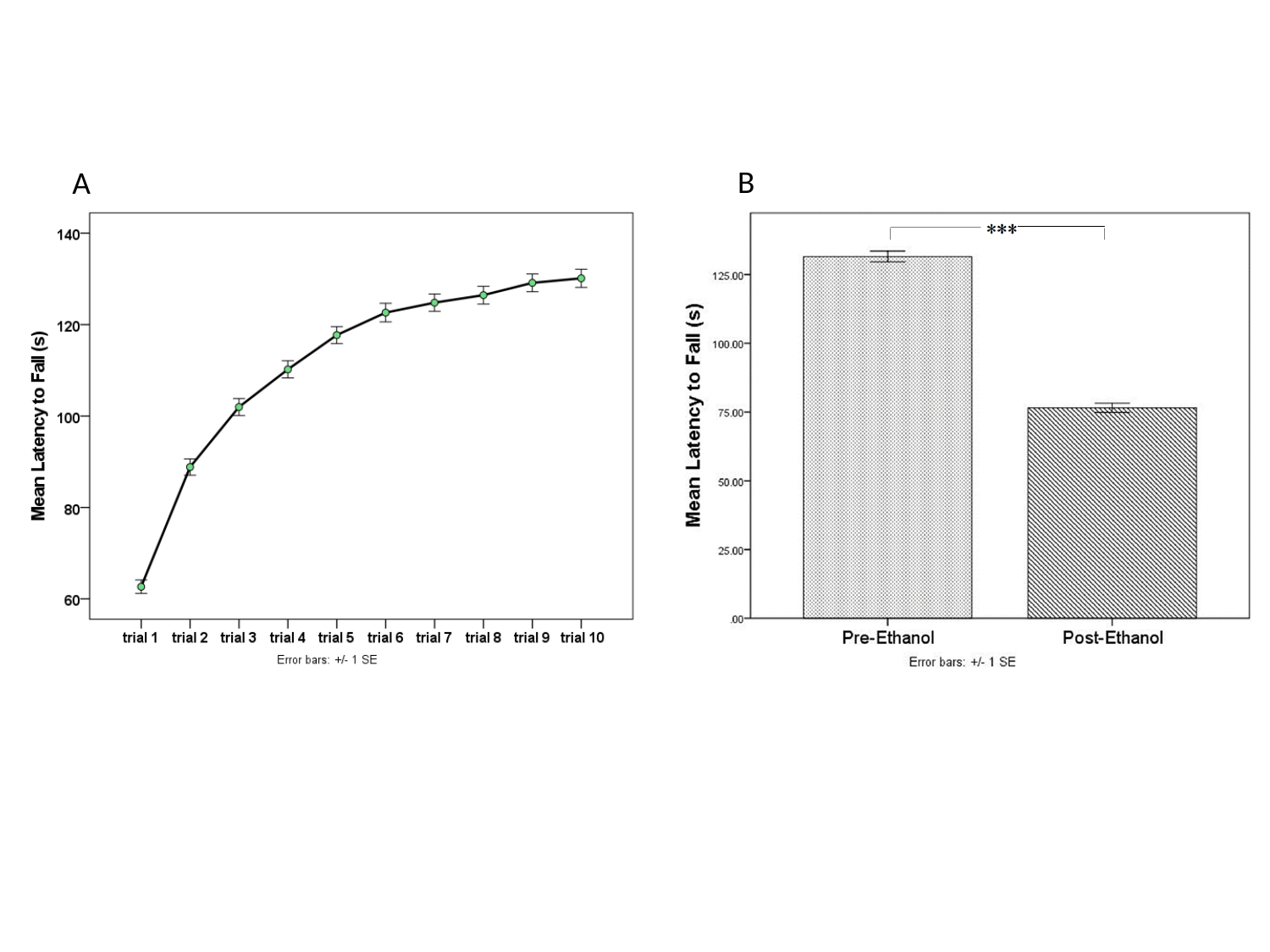

B
A

### Slide 6
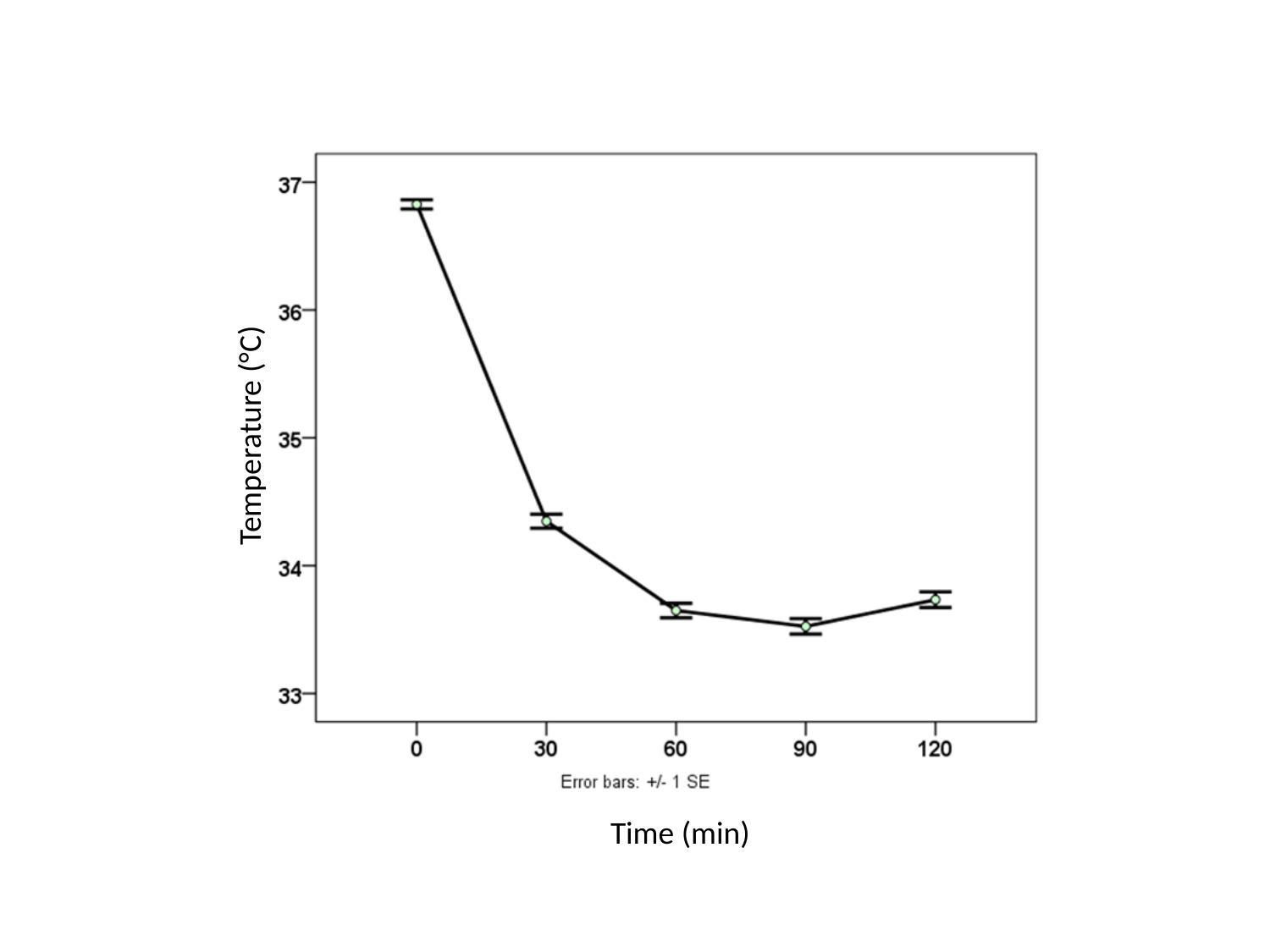

Temperature (°C)
Time (min)

### Slide 7
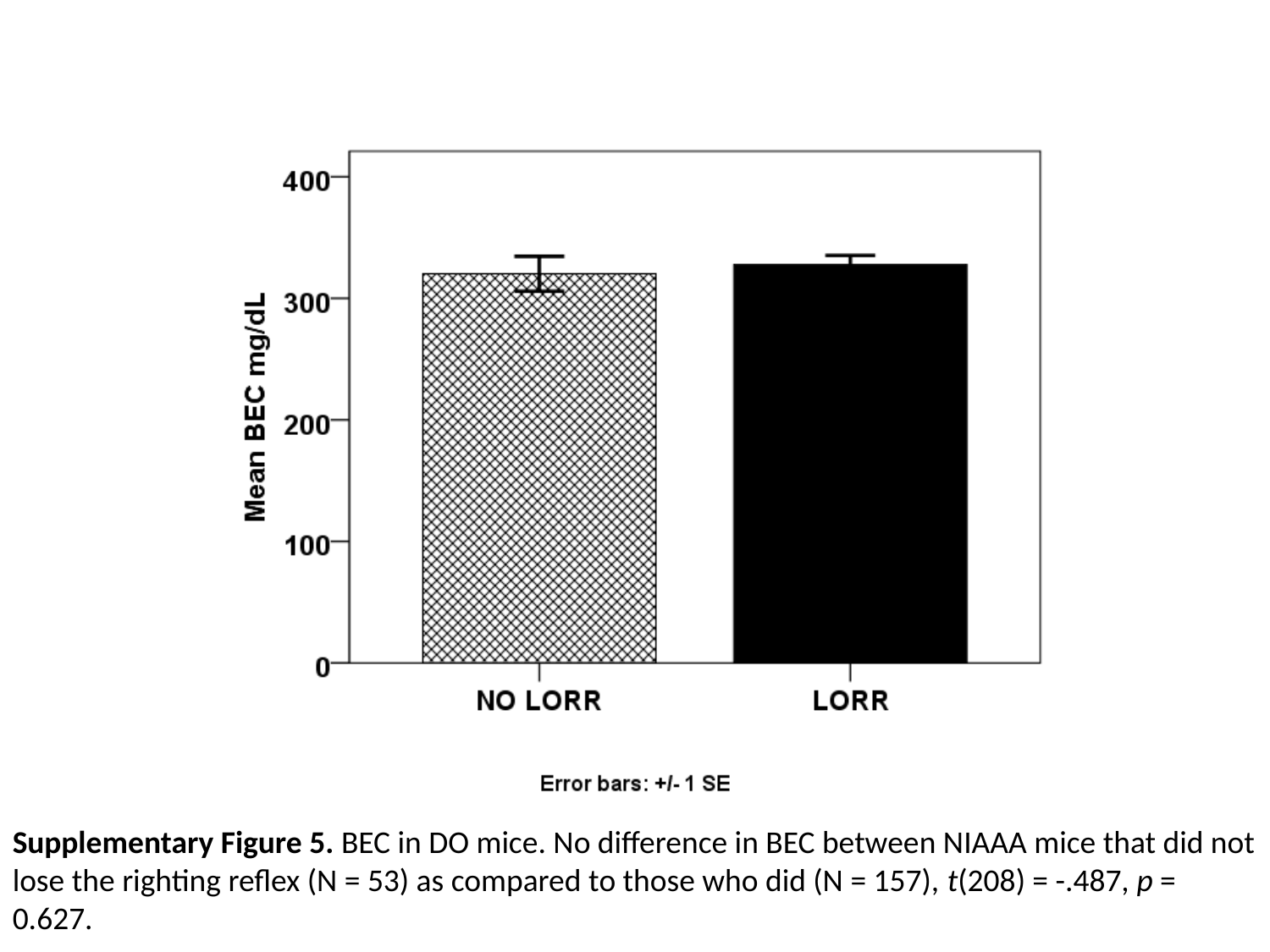

Supplementary Figure 5. BEC in DO mice. No difference in BEC between NIAAA mice that did not lose the righting reflex (N = 53) as compared to those who did (N = 157), t(208) = -.487, p = 0.627.

### Slide 8
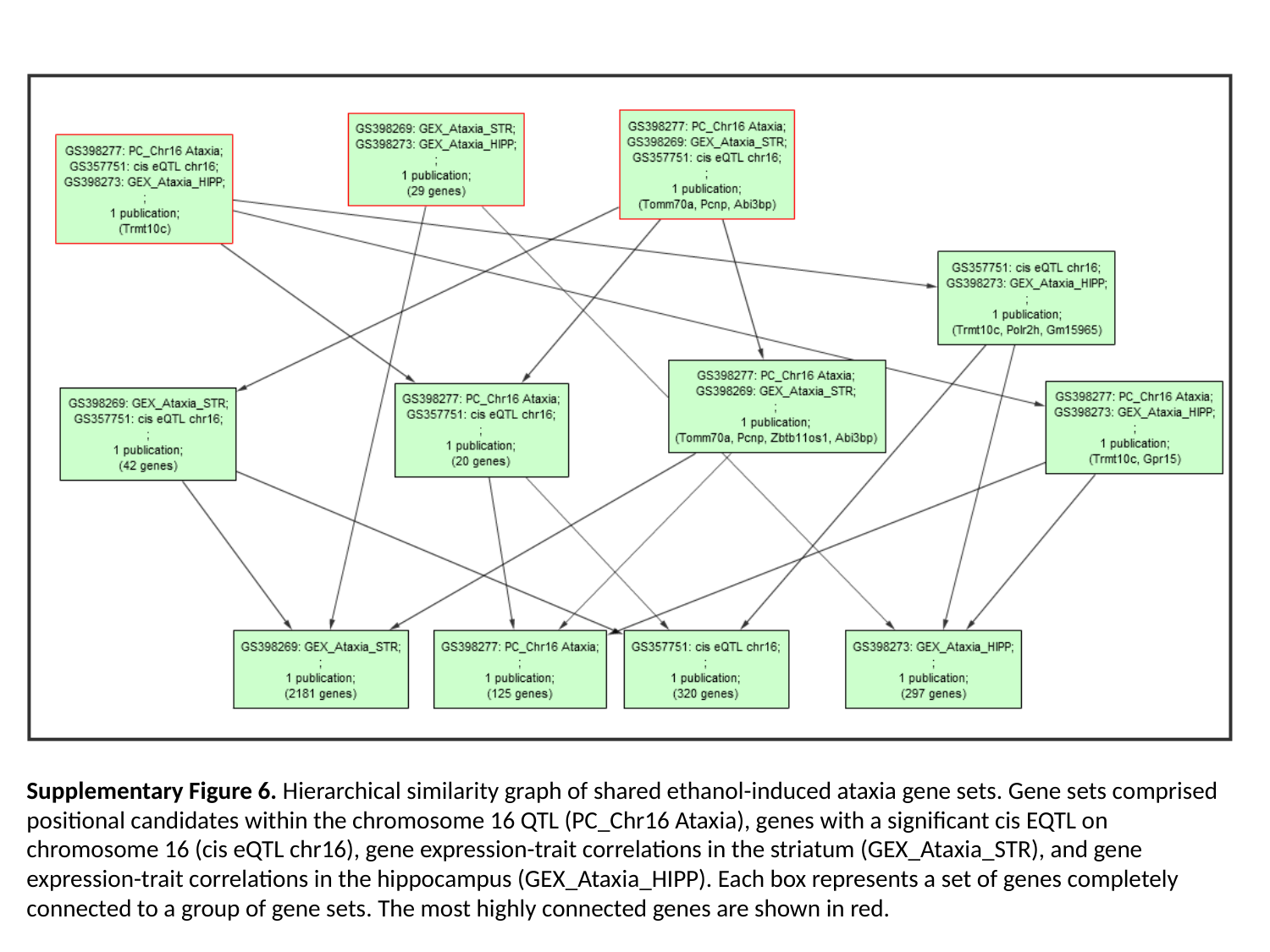

Supplementary Figure 6. Hierarchical similarity graph of shared ethanol-induced ataxia gene sets. Gene sets comprised positional candidates within the chromosome 16 QTL (PC_Chr16 Ataxia), genes with a significant cis EQTL on chromosome 16 (cis eQTL chr16), gene expression-trait correlations in the striatum (GEX_Ataxia_STR), and gene expression-trait correlations in the hippocampus (GEX_Ataxia_HIPP). Each box represents a set of genes completely connected to a group of gene sets. The most highly connected genes are shown in red.

### Slide 9
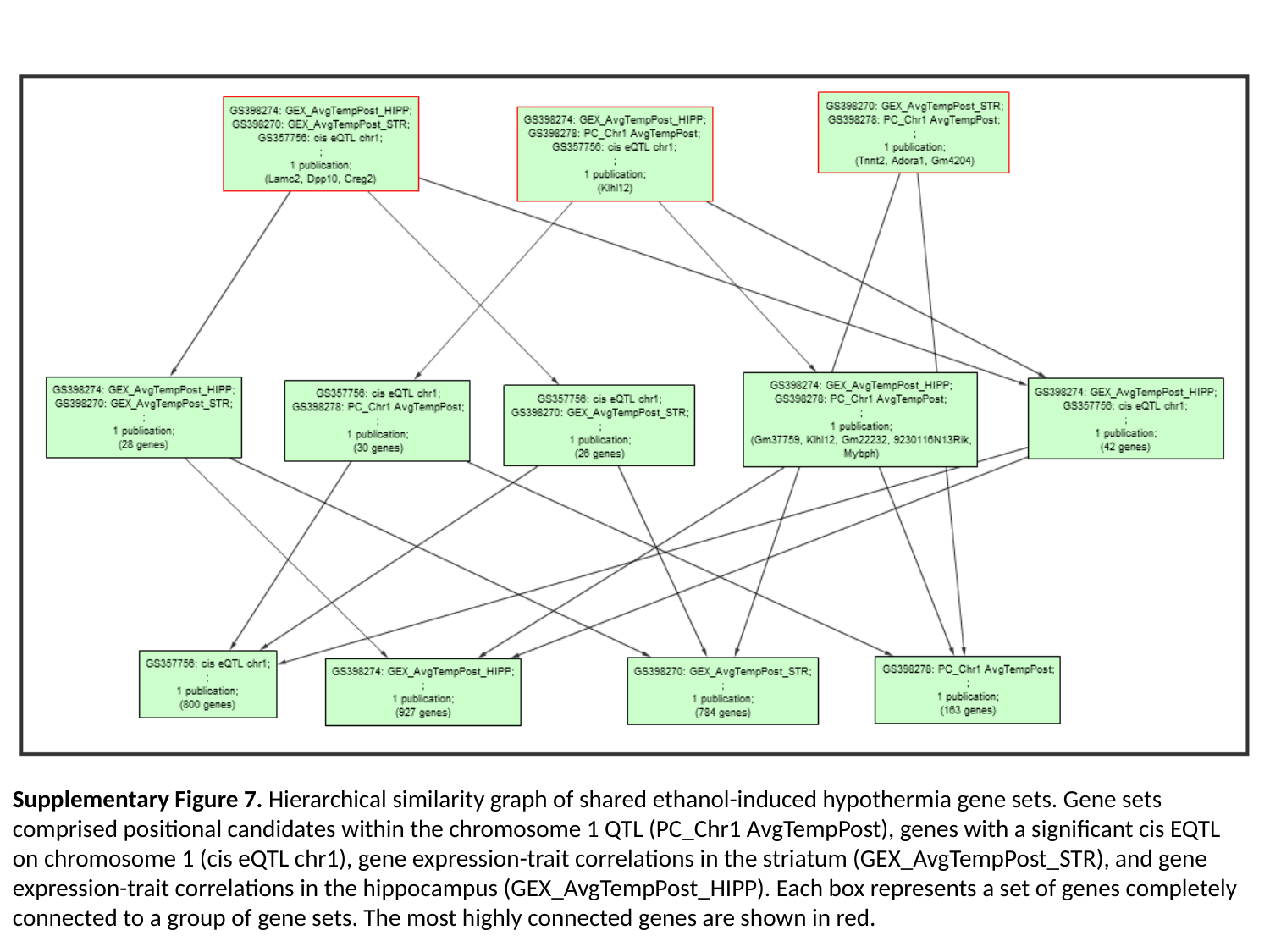

Supplementary Figure 7. Hierarchical similarity graph of shared ethanol-induced hypothermia gene sets. Gene sets comprised positional candidates within the chromosome 1 QTL (PC_Chr1 AvgTempPost), genes with a significant cis EQTL on chromosome 1 (cis eQTL chr1), gene expression-trait correlations in the striatum (GEX_AvgTempPost_STR), and gene expression-trait correlations in the hippocampus (GEX_AvgTempPost_HIPP). Each box represents a set of genes completely connected to a group of gene sets. The most highly connected genes are shown in red.

### Slide 10
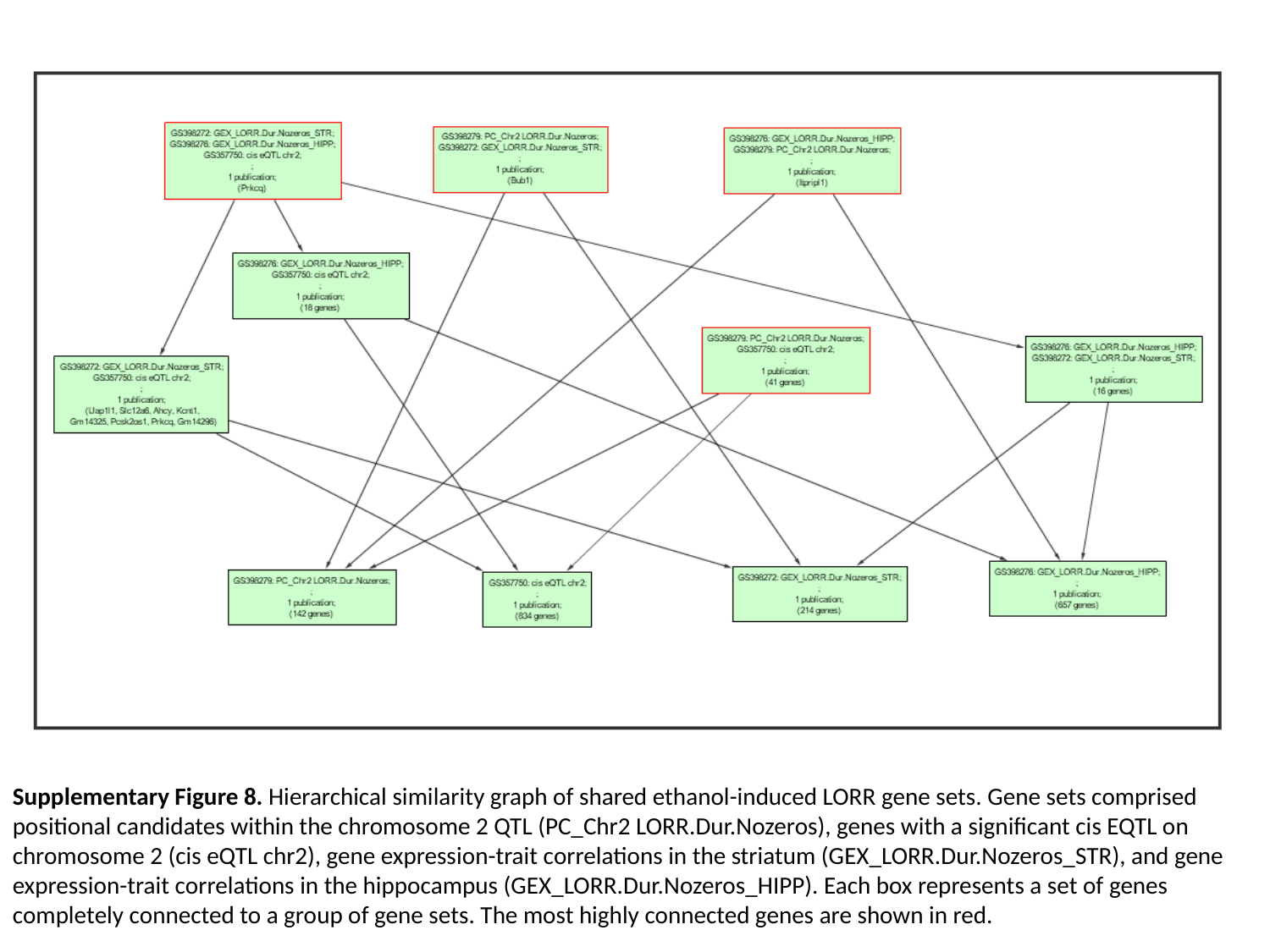

Supplementary Figure 8. Hierarchical similarity graph of shared ethanol-induced LORR gene sets. Gene sets comprised positional candidates within the chromosome 2 QTL (PC_Chr2 LORR.Dur.Nozeros), genes with a significant cis EQTL on chromosome 2 (cis eQTL chr2), gene expression-trait correlations in the striatum (GEX_LORR.Dur.Nozeros_STR), and gene expression-trait correlations in the hippocampus (GEX_LORR.Dur.Nozeros_HIPP). Each box represents a set of genes completely connected to a group of gene sets. The most highly connected genes are shown in red.
